# TIMAHAC: Streamlined Tandem IMAC-HILIC Workflow for Simultaneous and High-Throughput Plant Phosphoproteomics and N-glycoproteomics

**DOI:** 10.1101/2024.02.26.582001

**Authors:** Chin-Wen Chen, Pei-Yi Lin, Ying-Mi Lai, Miao-Hsia Lin, Shu-Yu Lin, Chuan-Chih Hsu

## Abstract

Protein post-translational modifications (PTMs) are crucial in plant cellular processes, particularly in protein folding and signal transduction. N-glycosylation and phosphorylation are notably significant PTMs, playing essential roles in regulating plant responses to environmental stimuli. However, current sequential enrichment methods for simultaneous analysis of phosphoproteome and N-glycoproteome are labor-intensive and time-consuming, limiting their throughput. Addressing this challenge, this study introduces a novel tandem S-Trap-IMAC-HILIC (S-Trap: suspension trapping; IMAC: immobilized metal ion affinity chromatography; HILIC: hydrophilic interaction chromatography) strategy, termed TIMAHAC, for simultaneous analysis of plant phosphoproteomics and N-glycoproteomics. This approach integrates IMAC and HILIC into a tandem tip format, streamlining the enrichment process of phosphopeptides and N-glycopeptides. The key innovation lies in the use of a unified buffer system and an optimized enrichment sequence to enhance efficiency and reproducibility. The applicability of TIMAHAC was demonstrated by analyzing the Arabidopsis phosphoproteome and N-glycoproteome in response to abscisic acid (ABA) treatment. Up to 1,954 N-glycopeptides and 11,255 phosphopeptides were identified from Arabidopsis, indicating its scalability for plant tissues. Notably, distinct perturbation patterns were observed in the phosphoproteome and N-glycoproteome, suggesting their unique contributions to ABA response. Our results reveal that TIMAHAC offers a comprehensive approach to studying complex regulatory mechanisms and PTM interplay in plant biology, paving the way for in-depth investigations into plant signaling networks.

## Introduction

Protein post-translational modifications (PTMs) are important molecular events in plant cells, involving protein folding and enzyme activation processes(1, 2). Among these, N-glycosylation and phosphorylation are particularly vital, as they regulate plant growth and response to environmental stresses(3–6). The complex landscape of PTMs is highlighted by multiple plant membrane proteins undergoing both phosphorylation and N-glycosylation, suggesting a potential interplay that influences protein function(7). Notable examples include receptor-like kinases such as BRI1(8, 9), FER(10, 11), and FLS2(12, 13), pivotal in plant hormone responses or innate immunity and are modified by both PTMs. However, a comprehensive understanding of the complex crosstalk between these modifications and their collective impact on kinase activity and downstream signaling remains elusive. A global profiling of plant phosphoproteome and N-glycoproteome over dynamic time courses could illuminate their synergistic roles in signal transductions and regulatory mechanisms.

Mass spectrometry (MS) has become a pivotal tool for systematically analyzing phosphorylation and N-glycosylation by identifying modification sites and unraveling N-glycan structures(14, 15). However, challenges such as the low abundance of modified peptides and N-glycosylation heterogeneity hinder direct MS detection of these peptides(16). Additionally, the ionization efficiency of phosphorylated and glycosylated peptides is typically lower due to the negative charge of the phosphate group or the hydrophilic nature of glycans (17). Various enrichment methods have been established to isolate these modified peptides prior to LC-MS/MS analysis(18, 19). For instance, immobilized metal affinity chromatography (IMAC) and metal oxide chromatography (MOAC) have been widely utilized for enriching phosphorylated peptides(20–22). Hydrophilic interaction chromatography (HILIC) stands out as a favored strategy for the enrichment of intact glycopeptides and N-glycoproteome profiling(23–26). Despite these developments, fully elucidating the interplay between phosphorylation and N-glycosylation remains challenging, due to the differing protocols and loading buffers required by these methods, which doubles the sample amount needed for parallel analysis and limits their use with low-input samples.

To overcome this bottleneck, two strategies have been developed for concurrent enrichment of both PTMs. The first strategy employs a single material to capture both PTMs, followed by sequential elution(27). For example, a dual-functional Ti^4+^-IMAC has been used for simultaneous enrichment of phosphopeptides, sialylated glycopeptides, neutral glycopeptides, and mannose-6-phosphate (M6P) glycopeptides(28). Stepwise elution was utilized to segregate N-glycopeptides with different glycan composition and phosphopeptides into multiple elution fractions.

Furthermore, a new material, epoxy-adenosine triphosphate (ATP)-Ti(IV)-IMAC, has been leveraged for the concurrent enrichment of phosphopeptides and N-glycopeptides, capitalizing on the high hydrophilicity of ATP and its metal ion chelation ability(29). The second strategy combines two methodologies for sequential enrichment of the two PTMs(30). Research groups proposed a workflow that integrates IMAC and anion exchange chromatography (MAX) in a two-step enrichment process(31, 32). Initially, Fe-IMAC is employed to capture phosphorylated peptides, followed by buffer exchange and subsequently N-glycopeptides enriched via MAX in the flow-through fraction. Despite these advancements, the workflows still necessitate multiple elution or buffer exchange steps, limiting the throughput for simultaneous phosphoproteome and N-glycoproteome analysis.

To streamline the multiple processes of plant phosphoproteomics, we developed a tandem suspension trapping (S-Trap)-IMAC workflow, replacing the conventional protein precipitation-based method typically used for eliminating small contaminates from plant lysate(33). This innovative approach allows for direct loading of digested peptides from S-Trap into a Fe-IMAC tip for phosphopeptide enrichment, thus circumventing buffer exchanges and reducing sample loss. Our results have presented enhanced sensitivity and robustness of this novel workflow when compared to the protein precipitation-based approach. This advancement now empowers large-scale studies into complex signaling pathways, even in sample-limited plant tissues. Inspired by these results, we explored integrating HILIC into this platform, enabling concurrent enrichment of phosphopeptides and N-glycopeptides, following by their separate elution for LC-MS/MS analysis.

We established a tandem S-Trap-IMAC-HILIC strategy for sequential enrichment of phosphopeptides and N-glycopeptides from plant samples. This involved systematic evaluation of the impact of TFA concentration on dual PTMs enrichment efficiency and optimization of the loading buffer for sequential enrichment. We determined the optimal sequence for PTMs enrichment, resolving how to prioritize phosphopeptide or N-glycopeptide enrichment based on PTMs identification coverage and quantification accuracy. Our findings partially elucidate why IMAC enrichment followed by HILIC yields better results than the reverse order. We demonstrated the applicability of this workflow in studying the crosstalk between phosphorylation and N-glycosylation in response to ABA treatment in Arabidopsis.

## Experimental Procedures

### Experimental Design and Statistical Rationale

Three technical replicates were conducted to calculate the coefficient of variation (CV) and mean for optimizing the sample loading buffer and benchmarking different enrichment strategies. Each sample was subjected to a single-shot LC-MS/MS analysis. The median identification number, mean intensity, and standard deviation (SD) of each condition was calculated and plotted in a bar chart. To examine the effects of 1 h ABA treatment on phosphoproteome and N-glycoproteome, four biological replicates of 10-day-old Arabidopsis seedlings were collected from two conditions. The seedlings without ABA treatment were used as controls, and 200 μg Arabidopsis lysate was used for the single-run phosphopeptide and N-glycopeptide enrichment experiments. A two-sample t-test was applied for differential phosphopeptides and N-glycopeptides analysis in terms of ABA treatment. An experimental design table, which includes all acquired files, is available in the PRIDE repository.

### Materials

Urea was purchased from Bio-Rad (Hercules, CA). Tris(2-carboxyethyl) phosphine hydrochloride (TCEP), 2-chloroacetamide (CAA), triethylammonium bicarbonate (TEAB), iron chloride, trifluoroacetic acid (TFA), ethylenediaminetetraacetic acid (EDTA), and sodium dodecyl sulfate (SDS) were purchased from Sigma-Aldrich (St. Louis, MO). Phosphoric acid (PA) was purchased from Honeywell Fluka (Charlotte, NC). Methanol (MeOH), formic acid (FA) and SeQuant ZIC-HILIC column were purchased from Merck (Darmstadt, Germany). Acetic acid (AA), acetonitrile (ACN), and ammonia phosphate (NH_4_H_2_PO_4_) were purchased from J.T.Baker (Radnor, PA). Ni-NTA silica beads were purchased from Qiagen (Hilden, Germany). Evotips were purchased from Evosep (Odense, Denmark). S-Trap micro columns were purchased from ProtiFi (Huntington, NY). Empore C8 extraction disks were purchased from 3M (St. Paul, MN). MS grade Lys-C (lysyl endopeptidase) was purchased from FUJIFILM Wako (Osaka, Japan). Sequencing-grade modified trypsin was purchased from Promega (Madison, WI). Water was obtained from a Millipore Milli-Q system (Bedford, MA).

### Plant Culture and ABA Treatment

Sterilized Arabidopsis Columbia-0 seeds were initiated by germination on 1/2 Murashige and Skoog (MS) medium at a temperature of 4°C for two days. For testing workflow conditions, the seedlings were subsequently cultivated vertically on plates for 14 days before being harvested. For experiments involving ABA treatment, the seedlings were cultivated vertically on plates for seven days, following which they were carefully transferred to a conical flask containing 1/2 MS medium and allowed to acclimatize for three days. Subsequently, the seedlings were subjected to treatment with or without 50 μM ABA for the specified durations as indicated.

### Protein Extraction and S-Trap Digestion

S-Trap digestion was executed following the S-trap-IMAC protocol(33). Arabidopsis seedlings were pulverized into a fine powder using liquid nitrogen and a mortar. The resulting powders were lysed in 8 M urea in 50 mM TEAB. The solution was transferred to a 1.7-mL tube and subjected to 10 cycles of sonication for 10 seconds each. Following this, the tube underwent centrifugation at 16,000 g and 4°C for 20 minutes, with the subsequent collection of the supernatant. The protein quantity was determined using the bicinchoninic acid assay (Thermo Fisher Scientific, Waltham, MA). A total of 100 μg of protein in the lysis buffer underwent reduction and alkylation using 10 mM TCEP and 40 mM CAA at 45°C for 15 minutes. Subsequently, the protein solution was supplemented with a final concentration of 5.5% (v/v) PA, followed by a six-fold volume of binding buffer (90% v/v MeOH in 100 mM TEAB). After gently vortexing the solution, it was loaded into an S-Trap micro column. The solution was then removed by centrifuging the column at 4,000 g for 1 minute. The column was subsequently subjected to three washes with 150 μL of binding buffer. Finally, 20 μL of a digestion solution containing 1 unit of Lys-C and 1 μg of trypsin in 50 mM TEAB was added to the column, which was then incubated at 47°C for 2 h. Each digested peptide was eluted from the S-Trap micro column using 120 μL of a solution composed of 1% (v/v) TFA in 80% (v/v) ACN, and these eluates were directly loaded into a Fe-IMAC tip, an HILIC tip, or tandem tips based on the experiment.

### Phosphopeptide Enrichment

Phosphopeptides were enriched following established methods with certain modifications(34, 35). To elucidate, an in-house IMAC tip was made by embedding a 20-μm polypropylene frit disk at the tip end, which was subsequently packed with Ni-NTA silica resin. The packed IMAC tip was inserted into a 2-mL Eppendorf tube. Initial removal of Ni^2+^ ions by adding 100 mM EDTA (200 g, 1 min). Subsequently, the tip was activated with 100 mM FeCl_3_ (200 g, 1 min) and equilibrated with 1% (v/v) TFA, 80% (v/v) ACN (200 g, 1 min). Tryptic peptides were eluted in 1% (v/v) TFA and 80% (v/v) ACN from S-Trap micro column and loaded onto the IMAC tip (200 g, 1 min). Sequential washing steps were executed utilizing 1% (v/v) TFA, 80% (v/v) ACN (200 g, 1 min), and 1% (v/v) acetic acid (pH 3.0) (200 g, 1 min) as the second washing step. The IMAC tip was then inserted into an activated Evotip. The captured phosphopeptides were eluted onto the Evotip using 200 mM NH_4_H_2_PO_4_ (200 g, 3 min), followed by placement of the Evotip into Evosep One LC system for LC-MS/MS analysis.

### N-glycopeptide Enrichment

The ZIC-HILIC silica resin was obtained from a SeQuant ZIC-HILIC column. The HILIC tip was made by embedding a C8 extraction disk at the tip’s end, and the resulting tip was inserted into a 2-mL Eppendorf tube. Activation of the C8 disk was performed by adding 1% (v/v) TFA, 80% (v/v) ACN (1000 g, 5 min). Once activated, the tip was subsequently packed with 20 mg of HILIC silica resin. The tip was washed using 1% (v/v) TFA (1000 g, 5 min), followed by 1% (v/v) TFA, 80% (v/v) ACN (1000 g, 5 min). Tryptic peptides were eluted in a solution composed of 1% (v/v) TFA and 80% (v/v) ACN from the S-Trap micro column and then loaded onto the HILIC tip (1000 g, 5 min). The tip was subjected to a series of three washes, utilizing 1% (v/v) TFA, 80% (v/v) ACN (1000 g, 5 min) each time. The HILIC tip was then inserted into an activated Evotip. The enriched N-glycopeptides were subsequently eluted onto the Evotip using 0.5% (v/v) FA (1000 g, 5 min), after which the Evotip was integrated into the Evosep One LC system for LC-MS/MS analysis.

### LC-MS/MS Analysis

The sample was loaded onto Evotips Pure and analyzed using the Evosep One LC system (EvoSep) coupled with a timsTOF HT mass spectrometer (Bruker Daltonics). Phosphopeptides and glycopeptides were separated analytically using the 30 SPD method with the Evosep Performance column (15 cm x 150 µm ID, 1.5 µm, EV1137; EvpSep) at 40 °C within a Captive Spray source (Bruker Daltonics), equipped with a 20 µm emitter (ZDV Sprayer 20, Bruker Daltonics).

Data acquisition on the timsTOF HT was performed using timsControl 4.0 (Bruker Daltonics) and operated in DDA-PASEF mode. The TIMS dimension was calibrated using the Agilent ESI LC/MS tuning mix with the following m/z values and 1/K0 values in positive mode: (622.0289, 0.9848 Vs/cm2), (922.0097, 1.1895 Vs/cm2), and (1221.9906, 1.3820 Vs/cm2). For phosphopeptide analysis, ten PASEF/MSMS scans were acquired per cycle. All spectra were acquired within an m/z range of 100 to 1,700 and an Ion Mobility (IM) of 0.6 to 1.6 V s/cm2. The accumulation and ramp time were set at 100 ms, the capillary voltage was maintained at 1400 V, and the values for ion mobility-dependent collision energy were linearly ramped from 20 eV at 0.60 V s/cm2 to 59 eV at 1.6 V s/cm2, remaining constant above or below. Singly charged precursors were excluded based on their position in the m/z-IM plane using a polygon shape, and precursor signals with an intensity threshold of 2,500 were selected for fragmentation. Precursors were isolated with a 2 Th window below m/z 700 and a 3 Th window above m/z 800. Additionally, they were actively excluded for 0.4 min when reaching a target intensity of 20,000.

For glycopeptides analysis, the m/z range was set 100 to 3,000. Stepping collision energy (CE) was applied for glycopeptide analysis for efficient glycopeptide dissociation. The stepping PASEF MS/MS frame consisted of two merged TIMS scans acquired for low and high CE profiles for glycopeptides. TIMS stepping was applied with two CEs: 32–64 eV was initially used, followed by 40–100 eV. Six PASEF MS/MS scans were triggered per cycle, with a target intensity of 40,000 for each individual PASEF precursor. The scan range was set between 0.68 and 1.66 V s/cm2.

### Data Processing

For phosphopeptides and N-glycopeptides identification, timsTOF raw files were converted into MGF format and subsequently searched against the Araport11 database (48,266 entries, release date: 2022-09-15) employing the Byonic search engine (version 4.3.4)(36) integrated within the Proteome Discoverer 2.5 (Thermo Fisher Scientific). Trypsin was selected as the digestion enzyme, permitting up to two missed cleavages. The static modification was set as carbamidomethyl (C), and variable modifications were set as oxidation (M), acetylation (protein N-term), phosphorylation (STY), and N-glycosylation. A precursor mass tolerance of 20 ppm and a fragment mass tolerance of 15 ppm were set. Identifications were filtered at 1% peptide-level and protein-level FDR. N-glycan modifications were searched against the N-glycan 52 plants database included in the Byonic software. N-glycopeptides (GPs) and glycopeptide-spectrum matches (GPSMs) were filtered using Byonic score >150, FDR 2D < 0.01, PEP 2D <0.05, and |log Prob| > 1.

### Data Analysis

N-Glycopeptides were exclusively categorized into five glycosylation-type categories based on the glycan composition identified: (1) truncated (1-2 HexNAc), (2) paucimannose (2 HexNAc and 1-3 Hex), (3) high mannose (2 HexNAc and > 3 Hex), (4) hybrid (3 HexNAc and 3-5 Hex), and complex (4 HexNAc and 3-5 Hex). Paucimannose, high mannose, hybrid, and complex categories may or may not contain fucose (Fuc) or xylose (Pent). Quantification of phosphopeptide was conducted using SpectroMine (version 4.2). PTM localization was set 0 and PEP.Label-Free Quant values were used for further quantitative analysis. Identified N-glycopeptides were extracted XIC intensity using Byos (version 5.2.31). Averaged intensity and CV values were calculated using the output tables of SpectroMine or Byos in Perseus (version 2.0.5.0)(37). Peptide collapse (version 1.4.4)(38) collapsed phosphopeptide output tables from SpectroMine, and set localization cutoff of 0.75 (class I sites). Quantifiable phosphorylation sites and glycopeptides were selected as identified in at least three replicates in at least one condition. The intensities were subsequently log2-transformed, and missing values were imputed using ImputeLCMD. Significant changes in phosphorylation sites and glycopeptides were identified using a two-sample t-test, with a *p*-value threshold of 0.05 and a minimum fold change (FC) of two. The significantly changed phosphorylation sites and glycopeptides were z-scored and used for hierarchical clustering analysis. Gene ontology enrichment analysis was executed via Fisher’s exact test, with parameters set to *p* < 0.001, and a FC > 2, utilizing the Panther database(39).

## Results

### Tandem S-Trap-IMAC-HILIC Workflow

In conventional sequential enrichment of phosphopeptides and N-glycopeptides, the utilization of distinct loading buffers necessitates multi-step processes. These steps, while necessary, often result in significant sample loss, adversely affecting detection sensitivity, reproducibility, and overall analysis throughput. To address these limitations, we develop a novel tandem S-Trap-IMAC-HILIC workflow, termed TIMAHAC, specifically tailored for the simultaneous analysis of plant phosphoproteomics and N-glycoproteomics (Fig. 1). The TIMAHAC approach innovatively utilizes a unified buffer system, serving dual purposes: peptide elution from the S-Trap and concurrent enrichment of phosphopeptides and N-glycopeptides. This integration is facilitated through a format of tandem S-Trap-IMAC-HILIC tip, which allows for simultaneous enrichment via simple centrifugation (supplemental Fig. S1A). Such a configuration effectively eliminates the need for multiple tube collections and sample transfers, thereby significantly reducing sample loss and variations typically encountered in sequential enrichment steps. Moreover, we have refined the procedure for loading the enriched phosphopeptides and N-glycopeptides onto an Evotip, followed by direct elution into the Evosep One system. This is achieved using a tandem arrangement of IMAC tip-Evotip and HILIC tip-Evotip (supplemental Fig. S1B and S1C). This advancement in our workflow negates the need for the conventional steps of peptide cleanup, buffer exchanges, tube collection and SpeedVac drying, streamlining the process and enhancing the efficiency of the overall protocol.

**Figure 1.**
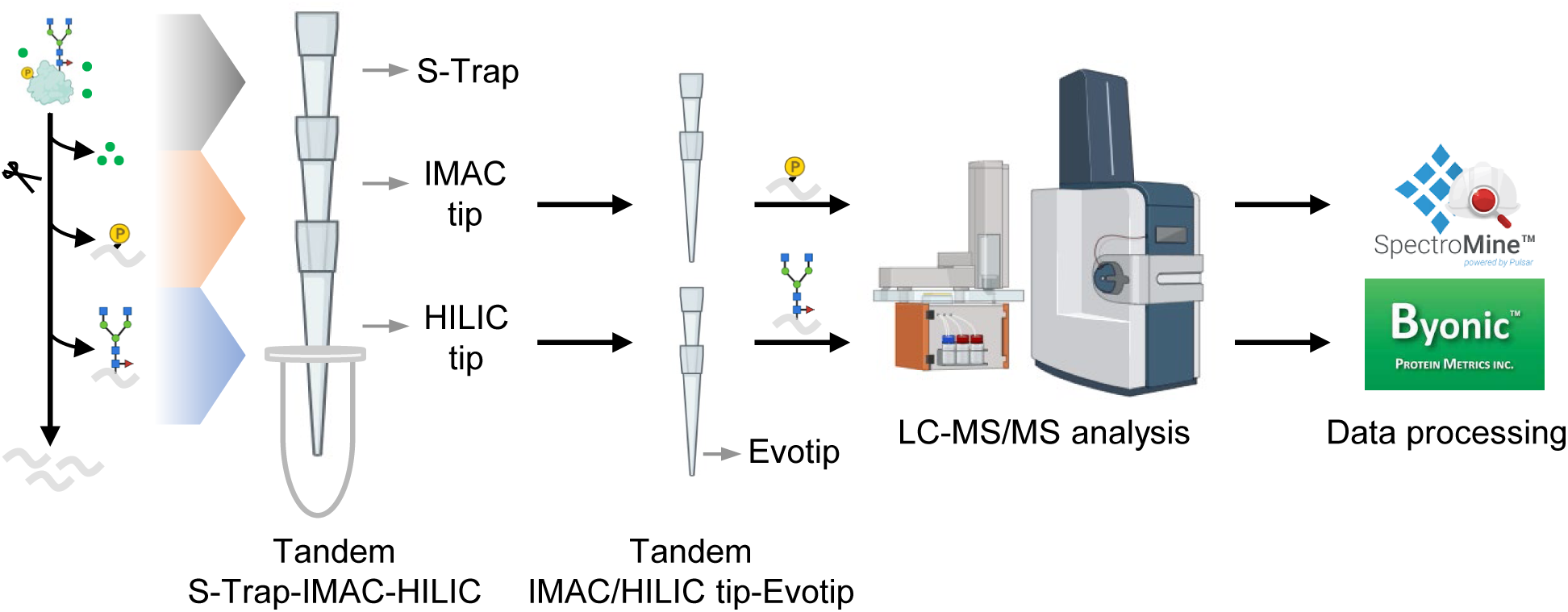
Experimental design of the TIMAHAC workflow for simultaneous analysis of plant phosphoproteomics and N-glycoproteomics. Contaminants removal and protein digestion are conducted within an S-Trap micro column. Phosphopeptides and N-glycopeptides are concurrently enriched through centrifugation using the tandem S-Trap-IMAC-HILIC strategy. The enriched peptides are directly loaded onto an Evotip and subsequently analyzed using an Evosep system coupled with a timsTOF HT mass spectrometer in DDA mode. Phosphoproteomics and N-glycoproteomics data are analyzed using SpectroMine and Byonic, respectively.

### Optimization of Loading Buffer for Tandem Tip-Based Enrichment

Optimization of the TFA concentration in the loading buffer is critical for enhancing the selectivity of phosphopeptides and N-glycopeptides(21, 24). This optimization is essential for enabling the utilization of a single loading buffer for the concurrent enrichment of phosphopeptides and N-glycopeptides in tandem tip-based format. To investigate how TFA concentration affects enrichment coverage and selectivity, we conducted experiments with four TFA concentrations (0.1%, 0.5%, 1%, and 2% (v/v)) in the IMAC and HILIC loading buffers, prepared in 80% ACN (supplemental Fig. S2A).

The evaluation of IMAC enrichment revealed that the identification of phosphorylated peptides and enrichment selectivity peaked at 0.5% TFA. This condition, however, did not significantly differ from the 1% TFA condition (Fig. 2A). A notable trend was observed in the identification of multiply phosphorylated peptides, which increased by approximate 39% when the TFA concentration was raised from 0.1% to 2%. However, there was a marked decline in monophosphorylated peptides, decreasing from 4800 to 3466, as TFA concentration exceeded 1%. This decrease is likely due to the incomplete deprotonation of the phosphate group on monophosphorylated peptides under low pH conditions. Additionally, the analysis of the accumulated extracted ion chromatography (XIC) area for identified phosphopeptides across the four TFA conditions indicated that 1% TFA provided the highest MS signals, balancing the signals from mono- or multiply phosphorylated peptides (supplemental Fig. S2B).

**Figure 2.**
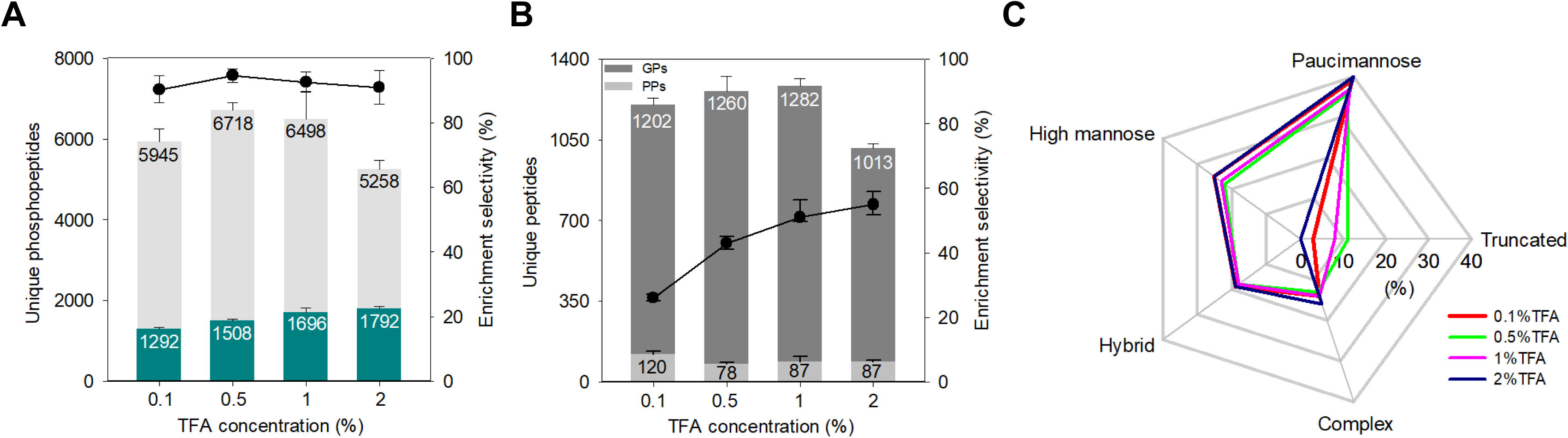
Evaluation of IMAC and HILIC enrichment efficiency at different TFA concentrations. A, number of identified phosphopeptides and multiply phosphorylated peptides by IMAC under four TFA concentration conditions. The light grey bar indicates the total number of identified phosphopeptides in each condition, while the dark teal bar represents the number of multiply phosphorylated peptides identified in each condition. The dots plot indicates the percentage of identified phosphopeptides relative to the total identified peptides (n = 3). Error bars, SD. B, number of identified N-glycopeptides and phosphopeptides using HILIC enrichment under different TFA concentration conditions. The dots plot illustrates the N-glycopeptide selectivity of HILIC in each condition (n = 3). Error bar, SD. C, radar plot summarizing the distribution of five glycan categorizations of identified unique glycoforms under four TFA concentration conditions. GPs, N-glycopeptides. PPs, phosphopeptides. SD, standard deviation.

We then assessed HILIC performance across the same TFA concentration range. The selectivity of HILIC improved from 26% to 55% as the TFA concentration increased from 0.1% to 2% (Fig. 2B). This enhancement is attributed to the ion-pairing effect of TFA, which neutralizes the charge on non-glycosylated peptides, thereby facilitating the separation of N-glycopeptides from non-glycopeptides during HILIC enrichment. The 1% TFA condition yielded the highest number of identified N-glycopeptides and GPSMs (supplemental Fig. S2C). However, an increase to 2% TFA leads to a significant reduction in the number of identified N-glycopeptides, likely due to the protonation of hydroxyl groups within the glycan moiety, which impedes hydrophilic partitioning within the HILIC stationary phase. The number of GPSMs and the XIC signals of identified N-glycopeptides remained relatively consistent across all TFA conditions (supplemental Fig. S2C and S2D). Further analysis of saccharide distribution in the glycan moieties of identified N-glycopeptides across the four TFA conditions revealed a distinct absence of N-glycopeptides with small glycan structures (< 6 saccharides) under the 2% TFA condition (supplemental Fig S3). This finding aligns with the glycan categorization distribution observed among the four TFA conditions (Fig. 2C). The 2% TFA condition exhibited an absence of truncated glycoforms and a relatively increase in high mannose and complex N-glycoforms compared to the 1% TFA condition.

In summary, the comprehensive evaluation of IMAC and HILIC performance under various TFA concentrations identified 1% TFA as the optimal concentration for the tandem tip-based workflow. This concentration effectively maximized the identification of phosphopeptides and N-glycopeptides, and it excelled in terms of MS signals and glycan group distribution. Thus, we selected an 80% ACN containing 1% TFA as the loading buffer for both IMAC and HILIC in all subsequent experiments.

### Determination of the Order of Sequential Phosphopeptides and N-glycopeptides Enrichment

Following the establishment of an optimal loading buffer for sequential enrichment of phosphopeptides and N-glycopeptides, our investigation focused on identifying the most effective enrichment order to ensure comprehensive coverage and to maintain reproducibility and quantitative accuracy of the plant phosphoproteome and N-glycoproteome. A key consideration was the potential loss of the second PTM during the enrichment steps of the first PTM. To explore the extent of sample loss during the enrichment of the first PTM, we evaluated two workflows: the tandem S-Trap-IMAC-HILC (IMAC-HILIC protocol) and S-Trap-HILIC-IMAC (HILIC-IMAC protocol) approaches (supplemental Fig. S4).

Initially, we compared phosphopeptide identification between the IMAC and HILIC-IMAC protocols to evaluate phosphopeptide loss during HILIC enrichment (Fig. 3A). As expected, phosphopeptide numbers decreased by approximately 9%—from 9560 to 8963—when passing through the HILIC tip. This reduction, particularly in monophosphorylated peptides, was likely due to retention within the HILIC stationary phase, as evidenced by phosphopeptide identification in the HILIC fraction (Fig. 2B). Notably, the decline in identified phosphopeptides after HILIC enrichment was primarily attributed to monophosphorylated peptides, with only a minor loss of less than 3% in multiply phosphorylated peptides (Fig. 3A). Next, we examined N-glycopeptide recovery in the HILIC and IMAC-HILIC protocols (Fig. 3B). Surprisingly, the IMAC-HILIC protocol showed a slight increase in identified N-glycopeptides, from 1124 to 1157, alongside an increase in HILIC selectivity from 59% to 73%. This suggests minimal N-glycopeptides loss during IMAC enrichment, corroborated by the absence of N-glycopeptides in the IMAC elution.

**Figure 3.**
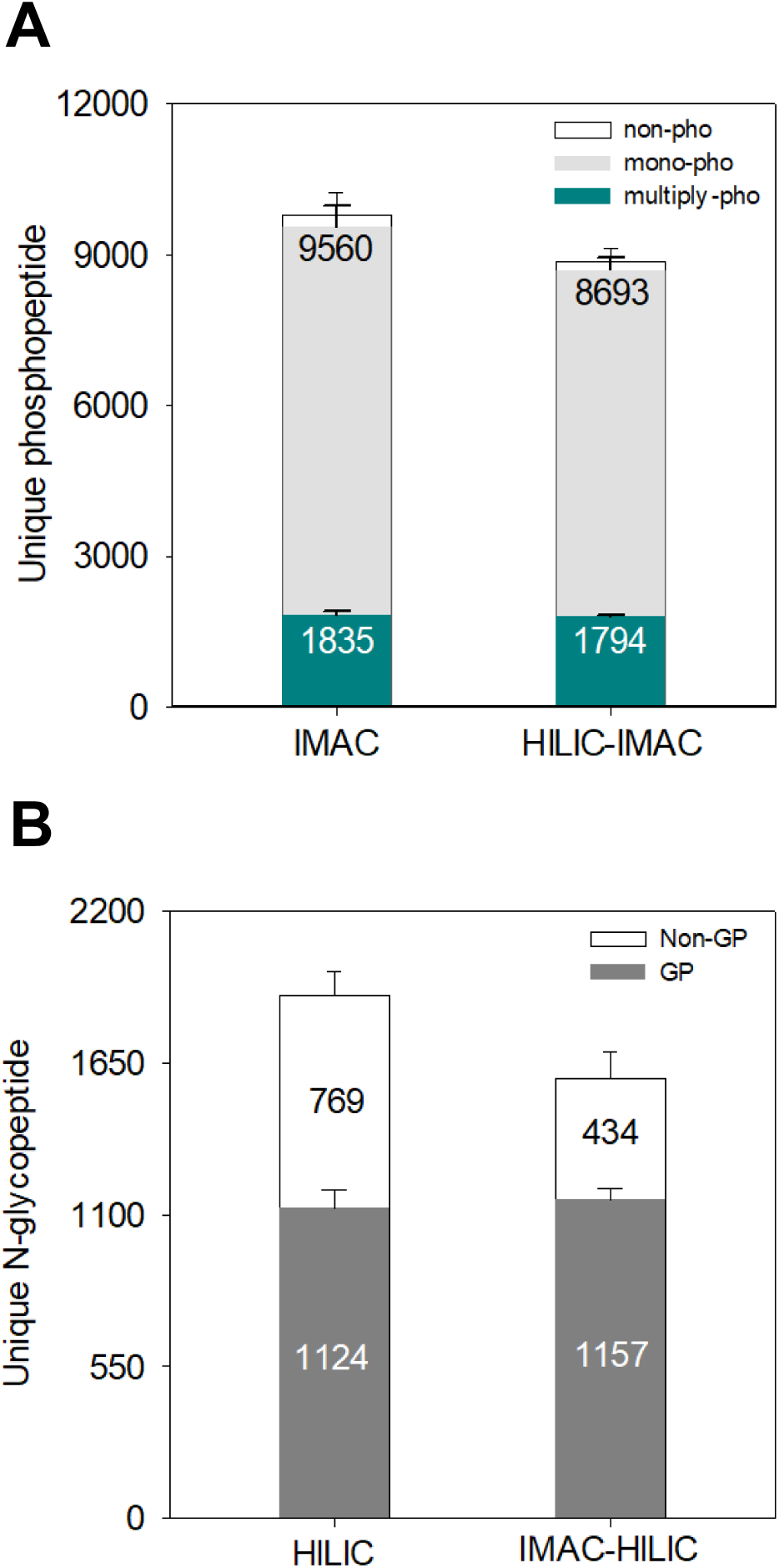
Benchmark of tandem IMAC-HILIC strategy against tandem HILIC-IMAC workflow. A, distribution of identified non-phosphopeptides, monophosphorylated peptides, and multiply phosphorylated peptides between the IMAC and HILIC-IMAC workflows (n = 3). Error bar, SD. B, distribution of identified non-glycopeptides and N-glycopeptides between the HILIC and IMAC-HILIC workflows (n = 3). Error bar, SD. Non-pho, non-phosphopeptide. Mono-pho, monophosphorylated peptide. Multiply-pho, multiply phosphorylated peptide. Non-GP, non-glycopeptide. GP, N-glycopeptide. SD, standard deviation.

To quantitatively evaluate phosphopeptide loss, we compared the total phosphopeptide XIC area between the IMAC and HILIC-IMAC protocols (Fig. 4A). A 28% decrease in MS intensity using the HILIC-IMAC protocol compared to the IMAC protocol was observed, aligning with the identification data indicating hydrophilic phosphopeptide loss during HILIC enrichment. Contrarily, N-glycopeptide analysis showed no significant difference in the XIC area of identified N-glycopeptides between protocols, suggesting that N-glycopeptides did not experience a significantly quantitative loss during IMAC enrichment (Fig. 4B).

**Figure 4.**
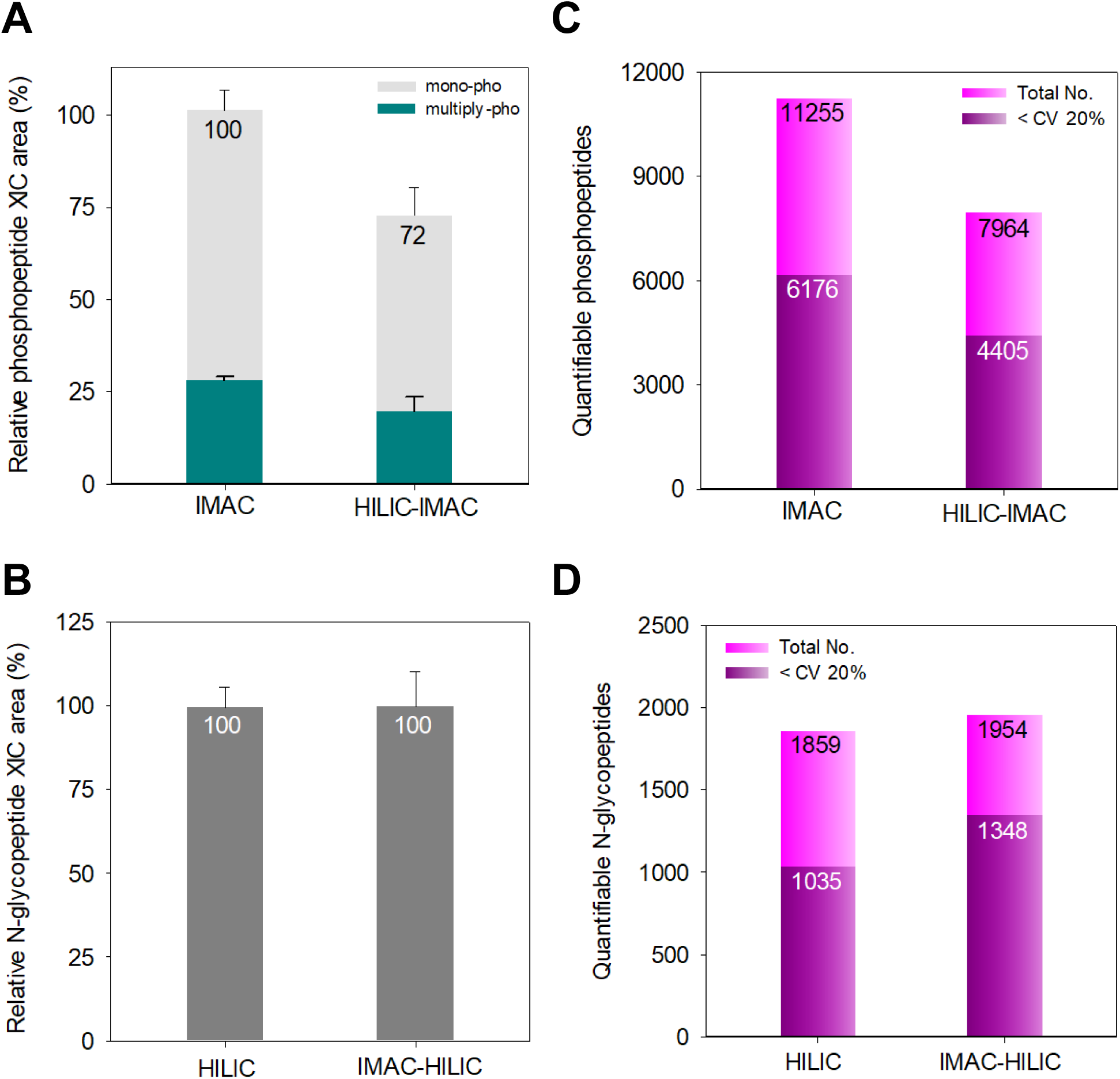
Performance assessment of MS signals and quantitative accuracy of in tandem HILIC-IMAC and IMAC-HILIC strategies. A, accumulated XIC area of identified monophosphorylated and multiply phosphorylated peptides compared between IMAC and HILIC-IMAC approaches (n = 3). Error bar, SD. B, accumulated XIC area of identified N-glycopeptides compared between HILIC and HILIC-IMAC approaches (n = 3). Error bar, SD. C, number of quantifiable phosphopeptides and phosphopeptides quantified with a CV below 20% compared between IMAC and HILIC-IMAC approaches. D, number of quantifiable N-glycopeptides and N-glycopeptides quantified with a CV below 20% compared between HILIC and IMAC-HILIC approaches. Mono-pho, monophosphorylated peptides. Multiply-pho, multiply phosphorylated peptides. Total No., total identified number. SD, standard deviation.

Furthermore, we assessed the quantitative reproducibility and accuracy of both protocols. The Pearson correlation coefficients between technical replicates, revealing no significant difference between the two protocols, ranging from 0.913 to 0.925 (supplemental Fig. S5A). The median CVs for phosphopeptide reproducibility were identical at 18 % (supplemental Fig. S5B). These results suggest that the order of phosphopeptides enrichment before or after HILIC enrichment does not substantially affect the reproducibility of phosphopeptide enrichment. Notably, the IMAC protocol identified more quantifiable phosphopeptides, defined as those identified in at least two out of three replicates and exhibiting CVs below 20%, compared to the HILIC-IMAC protocol (Fig. 4C).

Comparing the Pearson correlations and median CVs for identified N-glycopeptides between replicates of the HILIC and IMAC-HILIC protocols, the latter showed higher coefficient values (range from 0.961 to 0.975) and superior median CV values for quantifiable N-glycopeptides (14% versus 18%) (supplemental Fig. S6A and S6B). These results were reflected in the number and percentage of the N-glycopeptides with CVs less than 20% for the two protocols (Fig. 4D). While the numbers of quantifiable N-glycopeptides were comparable between the two protocols, it is noteworthy that the IMAC-HILIC protocol identified a higher percentage of N-glycopeptides with CVs less than 20% (69%, 1348) compared to the percentage in the HILIC protocol (56%, 1035). These data demonstrated that the reproducibility and quantification accuracy of identified N-glycopeptides was improved after IMAC enrichment.

Together, our study demonstrated that the TIMAHAC strategy, utilizing the S-Trap-IMAC-HILIC protocol, offers the optimal approach for concurrent analysis of both the plant phosphoproteome and N-glycoproteome.

### Enhancement of N-glycopeptide Selectivity in IMAC-HILIC Protocols

The observed increase in N-glycopeptide selectivity within the IMAC-HILIC protocol, compared to the direct HILIC protocol, prompted us to hypothesize that this improvement might be attributed to the reduced complexity of peptide mixtures following IMAC enrichment. Specifically, we hypothesized that certain non-glycopeptides, co-enriched in the HILIC process, might have been captured and removed by IMAC, thus enhancing HILIC selectivity. To test this hypothesis, we compared the overlap of non-glycopeptides identified in the second washing solution of the IMAC protocol, with those in the eluates of HILIC and IMAC-HILIC protocols (supplemental Fig. S7A). Our findings revealed a substantial proportion (43%, 462) of non-glycopeptides identified in the HILIC protocol were also present in the IMAC washing solution, compared to a significantly lower percentage (15%, 90) in the IMAC-HILIC protocol (Fig. 5A). This indicates that many non-glycopeptides co-enriched in the HILIC results were also captured by IMAC and subsequently eliminated during the washing step. This elucidates the reduced presence of non-glycopeptides after passing through the IMAC tip in the IMAC-HILIC protocol, thereby resulting in improved HILIC selectivity for N-glycopeptides (Fig. 3B).

**Figure 5.**
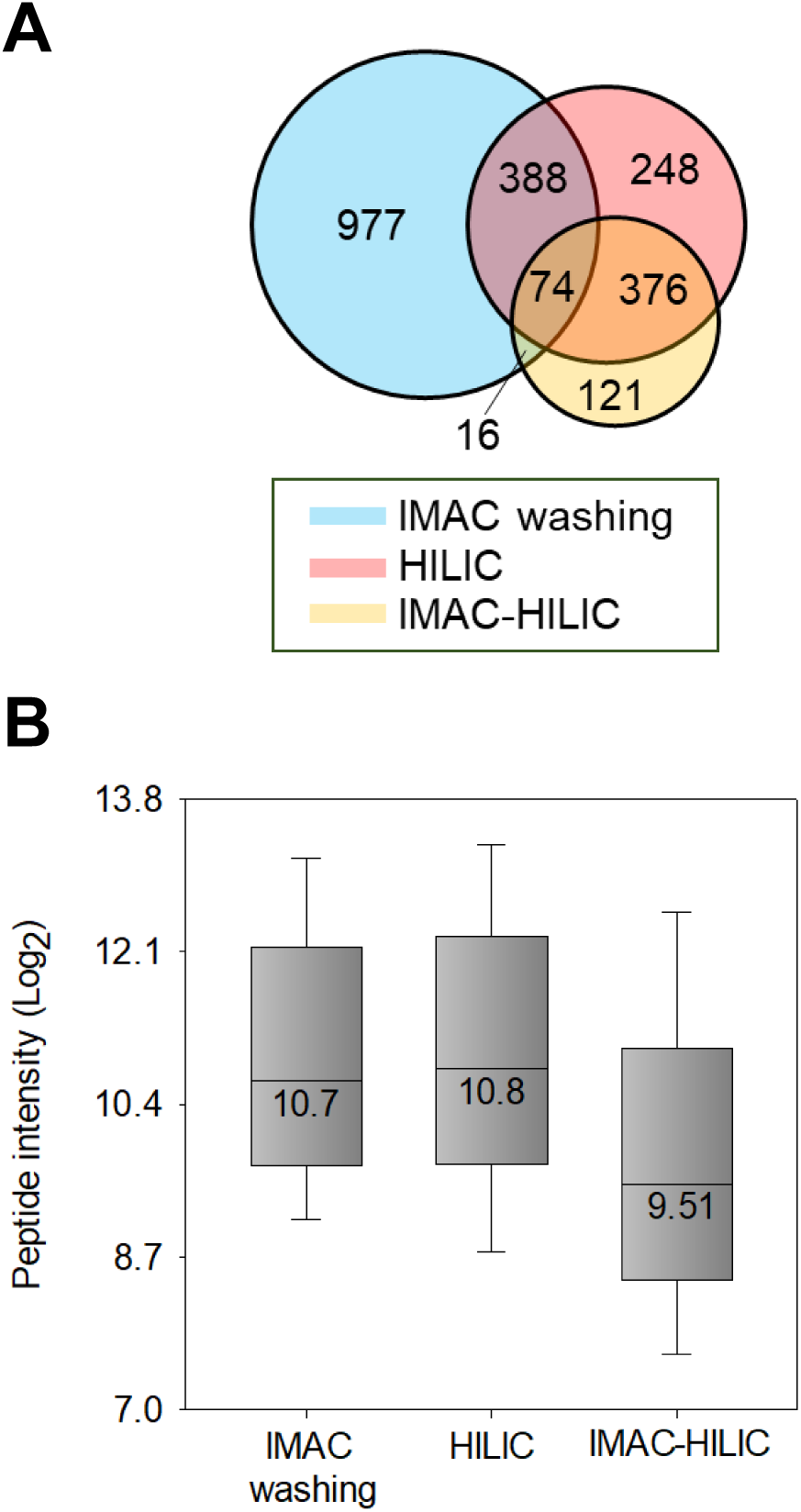
Comparison of the non-glycopeptide identification numbers and intensity medians in the IMAC washing, HILIC, and IMAC-HILIC results. A, Venn diagram representing the overlap of identified non-glycopeptides in three results. B, intensity distribution of the non-glycopeptides identified in all conditions. Boxes mark the first, median, and third quantiles, while whiskers mark the minimum/maximum value within 1.5 interquartile range. IMAC washing, the second step of IMAC washing.

Subsequently, we conducted a quantitative analysis of 450 non-glycopeptides identified in both HILIC and IMAC-HILIC protocols (Fig. 5A). This analysis revealed a 20% decrease in MS signals of non-glycopeptides in the IMAC-HILIC protocol (supplemental Fig. S7B), suggesting that the passage through IMAC effectively diminishes the abundance of these non-glycopeptides. This observation aligns with the comparison of the intensity distribution of 74 non-glycopeptides identified in all three conditions (Fig. 5B). Median intensities of non-glycopeptides identified in both the IMAC washing and HILIC were similar, whereas their median intensity in IMAC-HILIC was over 50% lower than in other conditions. Overall, our results indicate that the enhanced selectivity and quantification accuracy of N-glycopeptides observed in the IMAC-HILIC protocol could be attributed to the reduction in interference from hydrophilic non-glycopeptides, which are inevitably co-enriched in direct HILIC enrichment.

### In-Depth Analysis of ABA-Mediated N-Glycoproteome and Phosphoproteome in Arabidopsis

To demonstrate the scalability and applicability of the TIMAHAC method in the concurrent analysis of plant phosphoproteomics and N-glycoproteomics, we applied this approach to investigate the alterations in the ABA-dependent phosphoproteome and N-glycoproteome in Arabidopsis. We aimed to employ a rapid, highly sensitive approach to quantitative and accurate assessment of differential phosphorylation and N-glycosylation of key proteins in the ABA signaling pathway. To this end, we performed experiments on 1 h ABA-treated Arabidopsis seedlings (ABA) and untreated samples (control), with each condition replicated in four biological replicates. These samples were processed for phosphopeptide and N-glycopeptide using the TIMAHAC strategy, followed by analysis via timsTOF HT utilizing DDA-PASEF mode in 44 min. (supplemental Fig. S8A).

The TIMAHAC workflow exhibited high reproducibility in quantifying Arabidopsis phosphoproteomics and N-glycoproteomics, as indicated by median Pearson coefficients of 0.912 and 0.965, respectively (supplemental Fig. S8B and S8C). Single-shot analysis resulted in an average of 8,705 phosphopeptides and 1,221 N-glycopeptides from control samples and 9,357 phosphopeptides and 1,171 N-glycopeptides from ABA samples (Fig. 6A and 6B). We then assessed the perturbations in phosphorylation and N-glycosylation due to ABA treatment, observing that class I phosphosites were more induced (9.1%, 808) compared to the repressed phosphosites (5.7%, 506) (Fig. 6C and supplemental Table S1). In contrast, a significantly higher percentage of N-glycopeptides (10.1%, 231) were down-regulated in response to ABA treatment (Fig. 6D and supplemental Table S2). The distinct pattern highlights the unique contributions of these PTMs to the ABA response. Gene Ontology (GO) enrichment analysis further revealed unique GO biological processes between ABA-perturbated phosphoproteins and N-glycoproteins (supplemental Table S3). For example, there was a strong overrepresentation of phosphoproteins involved in organelle localization, mRNA splicing, vesicle-mediated transport, and protein phosphorylation are among the significantly ABA-modulated phosphoproteins (Fig. 6E). However, cell wall organization, polysaccharide metabolic process, developmental growth, and macromolecule biosynthetic process were overrepresented in ABA-perturbated N-glycoproteins (Fig. 6F). This divergence underscores the distinct biological functions of the phosphoproteome and N-glycoproteome in plant processes mediated by ABA.

**Figure 6.**
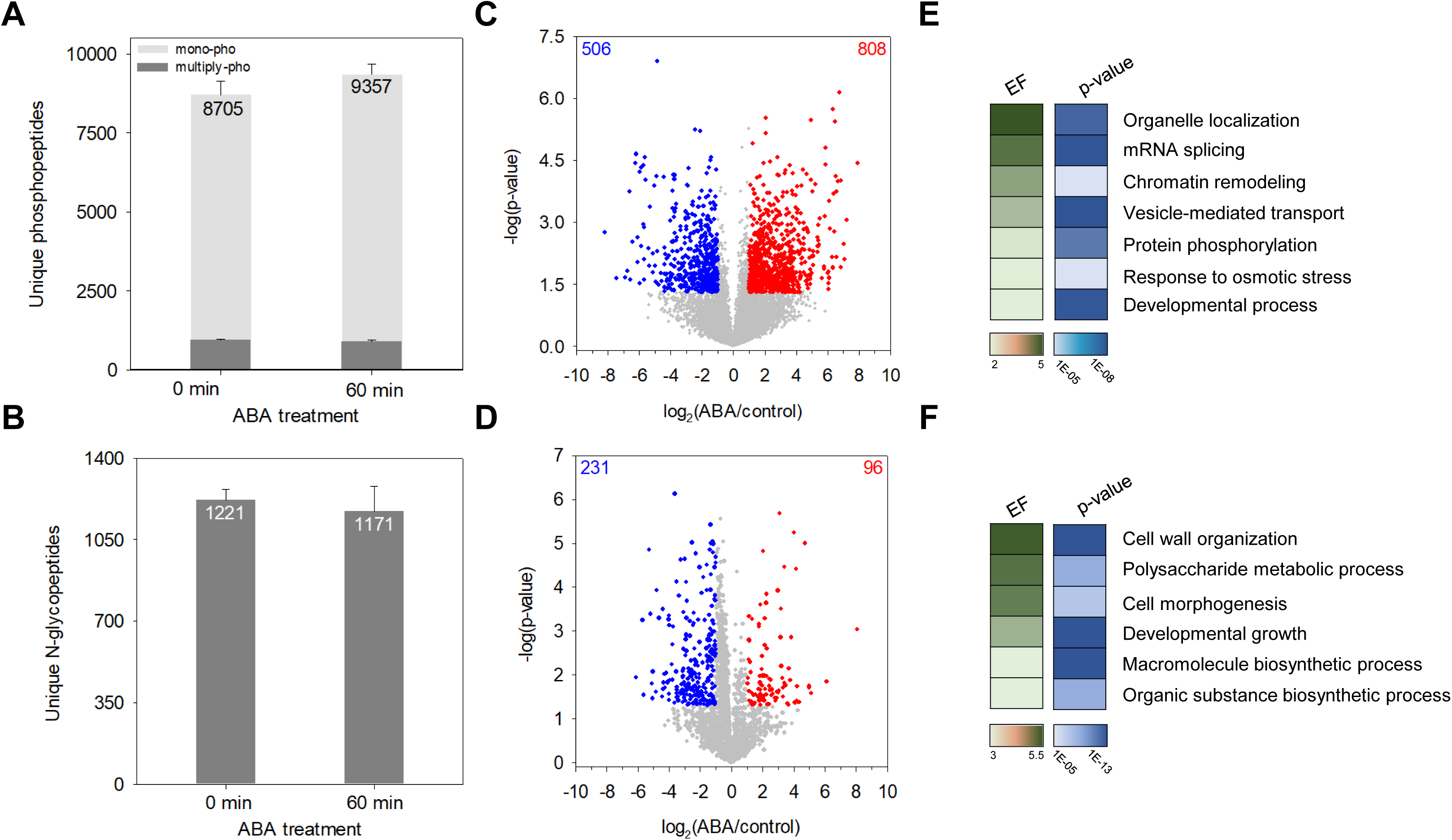
Investigation of ABA-mediated perturbation in Arabidopsis phosphoproteome and N-glycoproteome using the TIMAHAC workflow. A, number of identified phosphopeptides in control and ABA-treated Arabidopsis (n = 4). Error bars, SD. B, number of identified N-glycopeptides in control and ABA-treated Arabidopsis (n = 4). Error bars, SD. C, volcano plot of phosphorylation sites regulated upon 1 h of ABA treatment in Arabidopsis compared to untreated plants (*p* < 0.05, FC > 2 = red, FC < 2 = blue). D, volcano plot of N-glycopeptides regulated upon 1 h of ABA treatment in Arabidopsis compared to untreated plants (*p* < 0.05, FC > 2 = red, FC < 2 = blue). E, highlight of GOBP enriched by a Fisher’s exact test of phosphoproteins with significantly perturbated site upon ABA treatment (*p* < 0.001, EF > 2). F, highlight of GOBP enriched by a Fisher’s exact test of N-glycoproteins with significantly perturbated N-glycoform upon ABA treatment (*p* < 0.001, EF > 2). FC, fold change. EF, enrichment factor.

Further analysis focused on the impact of ABA on phosphorylation and N-glycosylation of kinases, revealing numerous phosphorylation sites on key cytosolic kinases involved in ABA signaling (supplemental Fig. S9A and S9B). For example, the phosphorylation of S177 on SnRK2.2, S172/S176 on SnRK2.3, and S175/T176 on SnRK2.6 were induced upon ABA treatment, and these phosphorylation sites are typically used to examine the activation of ABA signaling in Arabidopsis(40, 41). Several phosphorylation sites on casein kinase-like (CKL) proteins and mitogen-activated protein kinases (MAPKs) cascades were also induced by ABA(42, 43). Regarding kinases with perturbation of N-glycoform, 19 glycoforms corresponding to 15 N-glycosylated kinases were identified, primarily characterized as receptor-like kinases located in plasma membrane (supplemental Fig. S10A). A representative example is Feronia (FER), a crucial receptor-like kinase regulates plant growth and development(13); two of its N-glycoforms exhibited ABA-dependent changes. To investigate the potential cross-regulation between glycosylation and phosphorylation upon ABA treatment, we focused on the proteins modified by the two PTMs and perturbated by ABA. A total 9 proteins contain both ABA-perturbated phosphorylation and N-glycosylation sites (supplemental Fig. S10B). Remarkably, 4 out of the 9 proteins are kinases, suggesting the roles of kinases in the cross-regulation of the two PTMs in response to ABA. These include HDS-associated RLK1 (HAK1), impaired oomycete susceptibility 1 (IOS1), salt induced malectin-like domain-containing protein 1 (SIMP1), and barely any meristem 2 (BAM2). These proteins represent potential candidates for studying the interplay of phosphorylation and N-glycosylation in ABA signaling in plants. Together, this comprehensive analysis of the ABA-mediated phosphoproteome and N-glycoproteome in Arabidopsis, facilitated by the TIMAHAC method, provides valuable insights into plant stress responses and developmental pathways, highlighting the complex regulatory mechanisms underpinning these processes.

## Discussion

The conventional enrichment of both phosphopeptides and N-glycopeptides from plant samples can often be a time-consuming and labor-intensive process. We here introduced the TIMAHAC approach, a novel tandem S-Trap-IMAC-HILIC workflow, designed for simultaneous phosphoproteomics and N-glycopeptides in plant samples. Integrating these two enrichment methodologies, our aim was to overcome the inherent challenges of labor-intensive and sample loss-prone serial enrichment processes. Our innovative approach, employing a unified buffer for peptide elution and two PTMs enrichment, offers a streamlined workflow that abrogates the need for sample collection, additional peptide cleanup, buffer exchanges, and SpeedVac drying during sample preparation. As a result, it significantly reduces the total sample preparation time and enhances the throughput by enabling the simultaneous centrifugation of multiple tandem tips.

The TFA concentration in the loading buffer of the TIMAHAC approach plays a critical role in achieving comprehensive coverage and selectivity for both phosphopeptides and N-glycopeptides. Our experiments revealed that 1% TFA in the loading buffer was optimal for both IMAC and HILIC enrichment. It maximizes the number of identified phosphopeptides and N-glycopeptides while maintaining MS signals and their diverse glycan moieties. Further investigation into the order of sequential enrichment of phosphopeptides and N-glycopeptides was conducted to determine the most effective strategy while minimizing sample loss during the first PTM enrichment. A comparative analysis of both protocols indicated that HILIC-IMAC enrichment results in an approximate 30% loss of phosphopeptide signals. In contrast, N-glycopeptides did not experience significant losses during IMAC enrichment and demonstrated improved quantitative accuracy compared to direct HILIC enrichment. By comparing the non-glycopeptides existed in the IMAC washing and HILIC fraction, our data suggested that the improvement resulted from the reduction in both complexity and abundance of non-glycopeptides after IMAC enrichment. Based on these findings, we conclude that the IMAC-HILIC protocol is the more advantageous approach for concurrent enrichment of these two PTMs from plant samples.

It is noteworthy that in Arabidopsis, there was no significant co-enrichment observed for N-glycopeptides and phosphopeptides when using both IMAC and HILIC methods, respectively. This interesting result could be attributed to the unique nature of plant N-glycans and the Arabidopsis phosphoproteome. In contrast to human N-glycans, which contain sialic acid and M6P, plant N-glycans lack these components in their glycan structures(44). Sialylated and M6P-containing glycopeptides in human digests have been demonstrated to be co-enriched by IMAC materials through the electrostatic interactions between the negatively charged carboxylic groups or phosphate and the positively charged metal ions(28, 31). As a result, plant N-glycopeptides, lacking these negatively charged saccharides, are less likely to be captured during IMAC, thereby reducing their loss in the phosphopeptide enrichment process. Thus, no N-glycopeptides were co-identified in our IMAC eluates. In our HILIC results, we also observed less than 100 phosphopeptides identified with increasing TFA concentration beyond 0.1% (Fig. 2B). These numbers were significantly less than the numbers identified when using human cells as samples in HILIC(28). The discrepancy could be due to a lower abundance of acidic and highly hydrophilic phosphopeptides in the Arabidopsis phosphoproteome compared to that of human phosphoproteome, leading to a lower capture rate by the HILIC material. These reduced co-isolations of each PTM in these enrichment approaches thus facilitates a more distinct separation of the two PTMs when employing the TIMAHAC approach in plant systems.

In summary, the TIMAHAC strategy offers an optimal approach for the simultaneous analysis of plant phosphoproteomics and N-glycoproteomics. The unified buffer system, combined with careful TFA concentration optimization, ensures efficient and reproducible PTM enrichment. The IMAC-HILIC protocol demonstrates improved N-glycopeptide selectivity, providing an excellent platform for the comprehensive exploration of plant signaling networks influenced by these essential protein PTMs. The enhanced sensitivity, coverage, and quantification accuracy of this innovative approach, demonstrated through our study of ABA-dependent phosphoproteomic and N-glycoproteomic changes, pave the way for future research into the interplay between plant phosphorylation and N-glycosylation, and their roles in plant biology.

## Data availability

The mass spectrometry proteomics data have been deposited to the ProteomeXchange Consortium via the PRIDE partner repository(45) with the dataset identifier PXD047065. (Username; Password: BbQBUTEH)

## Supplemental data

This article contains supplemental data.

## Acknowledgements

This research was supported by grants (110-2311-B-001-043-MY2 and 112-2311-B-001-024-MY3) from the National Science and Technology Council. This work was also supported by Academia Sinica Core Facility and Innovative Instrument Project (AS-CII-111-209) from Academia Sinica. We thank Wen-Dar Lin from Bioinformatics Core of IPMB and Yu-Chun Chien from Kay-Hooi Khoo’s group for assisting data analysis. Our team would like to thank Chia-Feng Tsai for his helpful discussions. The research was performed in the Proteomics Core Lab, Institute of Plant and Microbial Biology, Academia Sinica.

## Author contributions

S.Y.L. and C.C.H. conceptualization; C.W.C., Y.M.L., and C.C.H methodology; C.W.C., P.Y.L., and Y.M.L. investigation; P.Y.L. resources; C.C.H. formal analysis; C.C.H. writing – original draft; M.H.L. and C.C.H. writing – review & editing; C.C.H. funding acquisition.

## Abbreviations

AA: acetic acid
ABA: abscisic acid
ACN: acetonitrile
CE: collision energy
CV: coefficient of variation
DDA: data-dependent acquisition
FA: formic acid
HILIC: hydrophilic interaction chromatography
IM: ion mobility
IMAC: immobilized metal ion affinity chromatography
LC: liquid chromatography
MS: mass spectrometry
PASEF: parallel accumulation-serial fragmentation
PTM: post-translational modification
SD: standard deviation
SPD: samples per day
S-Trap: suspension trapping
TFA: trifluoroacetic acid
TIMS: trapped ion mobility spectrometry
XIC: extracted ion chromatography

## Supplemental Data

**Supplemental Table S1**. Quantitative ABA-mediated phosphoproteomics data.

**Supplemental Table S2**. Quantitative ABA-mediated N-glycoproteomics data.

**Supplemental Table S3**. GO enrichment analysis.

## References

1. Friso, G., and van Wijk, K. J. (2015) Posttranslational Protein Modifications in Plant Metabolism. Plant Physiol 169, 1469–1487

2. Chong, L., Hsu, C. C., and Zhu, Y. (2022) Advances in mass spectrometry-based phosphoproteomics for elucidating abscisic acid signaling and plant responses to abiotic stress. J Exp Bot 73, 6547–6557

3. von Schaewen, A., Frank, J., and Koiwa, H. (2008) Role of complex N-glycans in plant stress tolerance. Plant Signal Behav 3, 871–873

4. Veit, C., Vavra, U., and Strasser, R. (2015) N-Glycosylation and plant cell growth. Methods Mol Biol 1242, 183–194

5. Wang, P., Zhao, Y., Li, Z., Hsu, C. C., Liu, X., Fu, L., Hou, Y. J., Du, Y., Xie, S., Zhang, C., Gao, J., Cao, M., Huang, X., Zhu, Y., Tang, K., Wang, X., Tao, W. A., Xiong, Y., and Zhu, J. K. (2018) Reciprocal Regulation of the TOR Kinase and ABA Receptor Balances Plant Growth and Stress Response. Mol Cell 69, 100–112 e106

6. Wang, W., Bai, M. Y., and Wang, Z. Y. (2014) The brassinosteroid signaling network-a paradigm of signal integration. Curr Opin Plant Biol 21, 147–153

7. Nagashima, Y., von Schaewen, A., and Koiwa, H. (2018) Function of N-glycosylation in plants. Plant Sci 274, 70–79

8. Jin, H., Yan, Z., Nam, K. H., and Li, J. (2007) Allele-specific suppression of a defective brassinosteroid receptor reveals a physiological role of UGGT in ER quality control. Mol Cell 26, 821–830

9. Kim, T. W., Guan, S., Burlingame, A. L., and Wang, Z. Y. (2011) The CDG1 kinase mediates brassinosteroid signal transduction from BRI1 receptor kinase to BSU1 phosphatase and GSK3-like kinase BIN2. Mol Cell 43, 561–571

10. Haruta, M., Sabat, G., Stecker, K., Minkoff, B. B., and Sussman, M. R. (2014) A peptide hormone and its receptor protein kinase regulate plant cell expansion. Science 343, 408–411

11. Lindner, H., Kessler, S. A., Muller, L. M., Shimosato-Asano, H., Boisson-Dernier, A., and Grossniklaus, U. (2015) TURAN and EVAN mediate pollen tube reception in Arabidopsis Synergids through protein glycosylation. PLoS Biol 13, e1002139

12. Gomez-Gomez, L., and Boller, T. (2000) FLS2: an LRR receptor-like kinase involved in the perception of the bacterial elicitor flagellin in Arabidopsis. Mol Cell 5, 1003–1011

13. Duan, Q., Kita, D., Li, C., Cheung, A. Y., and Wu, H. M. (2010) FERONIA receptor-like kinase regulates RHO GTPase signaling of root hair development. Proc Natl Acad Sci U S A 107, 17821–17826

14. Hsu, C. C., Zhu, Y., Arrington, J. V., Paez, J. S., Wang, P., Zhu, P., Chen, I. H., Zhu, J. K., and Tao, W. A. (2018) Universal Plant Phosphoproteomics Workflow and Its Application to Tomato Signaling in Response to Cold Stress. Mol Cell Proteomics 17, 2068–2080

15. Xu, S. L., Medzihradszky, K. F., Wang, Z. Y., Burlingame, A. L., and Chalkley, R. J. (2016) N-Glycopeptide Profiling in Arabidopsis Inflorescence. Mol Cell Proteomics 15, 2048–2054

16. Huang, J., Wang, F., Ye, M., and Zou, H. (2014) Enrichment and separation techniques for large-scale proteomics analysis of the protein post-translational modifications. J Chromatogr A 1372C, 1–17

17. Stavenhagen, K., Hinneburg, H., Thaysen-Andersen, M., Hartmann, L., Varon Silva, D., Fuchser, J., Kaspar, S., Rapp, E., Seeberger, P. H., and Kolarich, D. (2013) Quantitative mapping of glycoprotein micro-heterogeneity and macro-heterogeneity: an evaluation of mass spectrometry signal strengths using synthetic peptides and glycopeptides. J Mass Spectrom 48, i

18. Riley, N. M., Bertozzi, C. R., and Pitteri, S. J. (2021) A Pragmatic Guide to Enrichment Strategies for Mass Spectrometry-Based Glycoproteomics. Mol Cell Proteomics 20, 100029

19. Low, T. Y., Mohtar, M. A., Lee, P. Y., Omar, N., Zhou, H., and Ye, M. (2021) Widening the Bottleneck of Phosphoproteomics: Evolving Strategies for Phosphopeptide Enrichment. Mass Spectrom Rev 40, 309–333

20. Sugiyama, N., Masuda, T., Shinoda, K., Nakamura, A., Tomita, M., and Ishihama, Y. (2007) Phosphopeptide enrichment by aliphatic hydroxy acid-modified metal oxide chromatography for nano-LC-MS/MS in proteomics applications. Mol Cell Proteomics 6, 1103–1109

21. Tsai, C. F., Wang, Y. T., Chen, Y. R., Lai, C. Y., Lin, P. Y., Pan, K. T., Chen, J. Y., Khoo, K. H., and Chen, Y. J. (2008) Immobilized metal affinity chromatography revisited: pH/acid control toward high selectivity in phosphoproteomics. J Proteome Res 7, 4058–4069

22. Zhou, H., Ye, M., Dong, J., Han, G., Jiang, X., Wu, R., and Zou, H. (2008) Specific phosphopeptide enrichment with immobilized titanium ion affinity chromatography adsorbent for phosphoproteome analysis. J Proteome Res 7, 3957–3967

23. Hagglund, P., Bunkenborg, J., Elortza, F., Jensen, O. N., and Roepstorff, P. (2004) A new strategy for identification of N-glycosylated proteins and unambiguous assignment of their glycosylation sites using HILIC enrichment and partial deglycosylation. J Proteome Res 3, 556–566

24. Mysling, S., Palmisano, G., Hojrup, P., and Thaysen-Andersen, M. (2010) Utilizing ion-pairing hydrophilic interaction chromatography solid phase extraction for efficient glycopeptide enrichment in glycoproteomics. Anal Chem 82, 5598–5609

25. Jandera, P. (2011) Stationary and mobile phases in hydrophilic interaction chromatography: a review. Anal Chim Acta 692, 1–25

26. Chen, Y. J., Yen, T. C., Lin, Y. H., Chen, Y. L., Khoo, K. H., and Chen, Y. J. (2021) ZIC-cHILIC-Based StageTip for Simultaneous Glycopeptide Enrichment and Fractionation toward Large-Scale N-Sialoglycoproteomics. Anal Chem 93, 15931–15940

27. Huang, J., Dong, J., Shi, X., Chen, Z., Cui, Y., Liu, X., Ye, M., and Li, L. (2019) Dual-Functional Titanium(IV) Immobilized Metal Affinity Chromatography Approach for Enabling Large-Scale Profiling of Protein Mannose-6-Phosphate Glycosylation and Revealing Its Predominant Substrates. Anal Chem 91, 11589–11597

28. Huang, J., Liu, X., Wang, D., Cui, Y., Shi, X., Dong, J., Ye, M., and Li, L. (2021) Dual-Functional Ti(IV)-IMAC Material Enables Simultaneous Enrichment and Separation of Diverse Glycopeptides and Phosphopeptides. Anal Chem 93, 8568–8576

29. Wang, D., Huang, J., Zhang, H., Ma, M., Xu, M., Cui, Y., Shi, X., and Li, L. (2023) ATP-Coated Dual-Functionalized Titanium(IV) IMAC Material for Simultaneous Enrichment and Separation of Glycopeptides and Phosphopeptides. J Proteome Res 22, 2044–2054

30. Andaluz Aguilar, H., Iliuk, A. B., Chen, I. H., and Tao, W. A. (2020) Sequential phosphoproteomics and N-glycoproteomics of plasma-derived extracellular vesicles. Nat Protoc 15, 161–180

31. Cho, K. C., Chen, L., Hu, Y., Schnaubelt, M., and Zhang, H. (2019) Developing Workflow for Simultaneous Analyses of Phosphopeptides and Glycopeptides. ACS Chem Biol 14, 58–66

32. Zhou, Y., Lih, T. M., Yang, G., Chen, S. Y., Chen, L., Chan, D. W., Zhang, H., and Li, Q. K. (2020) An Integrated Workflow for Global, Glyco-, and Phospho-proteomic Analysis of Tumor Tissues. Anal Chem 92, 1842–1849

33. Chen, C. W., Tsai, C. F., Lin, M. H., Lin, S. Y., and Hsu, C. C. (2023) Suspension Trapping-Based Sample Preparation Workflow for In-Depth Plant Phosphoproteomics. Anal Chem 95, 12232–12239

34. Tsai, C. F., Hsu, C. C., Hung, J. N., Wang, Y. T., Choong, W. K., Zeng, M. Y., Lin, P. Y., Hong, R. W., Sung, T. Y., and Chen, Y. J. (2014) Sequential phosphoproteomic enrichment through complementary metal-directed immobilized metal ion affinity chromatography. Anal Chem 86, 685–693

35. Tsai, C. F., Wang, Y. T., Hsu, C. C., Kitata, R. B., Chu, R. K., Velickovic, M., Zhao, R., Williams, S. M., Chrisler, W. B., Jorgensen, M. L., Moore, R. J., Zhu, Y., Rodland, K. D., Smith, R. D., Wasserfall, C. H., Shi, T., and Liu, T. (2023) A streamlined tandem tip-based workflow for sensitive nanoscale phosphoproteomics. Commun Biol 6, 70

36. Bern, M., Kil, Y. J., and Becker, C. (2012) Byonic: advanced peptide and protein identification software. Curr Protoc Bioinformatics Chapter 13, 13 20 11–13 20 14

37. Tyanova, S., Temu, T., Sinitcyn, P., Carlson, A., Hein, M. Y., Geiger, T., Mann, M., and Cox, J. (2016) The Perseus computational platform for comprehensive analysis of (prote)omics data. Nat Methods 13, 731–740

38. Bekker-Jensen, D. B., Bernhardt, O. M., Hogrebe, A., Martinez-Val, A., Verbeke, L., Gandhi, T., Kelstrup, C. D., Reiter, L., and Olsen, J. V. (2020) Rapid and site-specific deep phosphoproteome profiling by data-independent acquisition without the need for spectral libraries. Nat Commun 11, 787

39. Thomas, P. D., Ebert, D., Muruganujan, A., Mushayahama, T., Albou, L. P., and Mi, H. (2022) PANTHER: Making genome-scale phylogenetics accessible to all. Protein Sci 31, 8–22

40. Wang, P., Xue, L., Batelli, G., Lee, S., Hou, Y. J., Van Oosten, M. J., Zhang, H., Tao, W. A., and Zhu, J. K. (2013) Quantitative phosphoproteomics identifies SnRK2 protein kinase substrates and reveals the effectors of abscisic acid action. Proc Natl Acad Sci U S A 110, 11205–11210

41. Lin, Z., Li, Y., Zhang, Z., Liu, X., Hsu, C. C., Du, Y., Sang, T., Zhu, C., Wang, Y., Satheesh, V., Pratibha, P., Zhao, Y., Song, C. P., Tao, W. A., Zhu, J. K., and Wang, P. (2020) A RAF-SnRK2 kinase cascade mediates early osmotic stress signaling in higher plants. Nat Commun 11, 613

42. Wang, P., Hsu, C. C., Du, Y., Zhu, P., Zhao, C., Fu, X., Zhang, C., Paez, J. S., Macho, A. P., Tao, W. A., and Zhu, J. K. (2020) Mapping proteome-wide targets of protein kinases in plant stress responses. Proc Natl Acad Sci U S A 117, 3270–3280

43. Zhang, M., and Zhang, S. (2022) Mitogen-activated protein kinase cascades in plant signaling. J Integr Plant Biol 64, 301–341

44. Strasser, R. (2016) Plant protein glycosylation. Glycobiology 26, 926–939

45. Perez-Riverol, Y., Bai, J., Bandla, C., Garcia-Seisdedos, D., Hewapathirana, S., Kamatchinathan, S., Kundu, D. J., Prakash, A., Frericks-Zipper, A., Eisenacher, M., Walzer, M., Wang, S., Brazma, A., and Vizcaino, J. A. (2022) The PRIDE database resources in 2022: a hub for mass spectrometry-based proteomics evidences. Nucleic Acids Res 50, D543–D552

