## Supplemental_figure_S1-S10 for "TIMAHAC: Streamlined Tandem IMAC-HILIC Workflow for Simultaneous and High-Throughput Plant Phosphoproteomics and N-glycoproteomics"

**TIMAHAC: Streamlined Tandem IMAC-HILIC Workflow for Simultaneous and High-Throughput Plant Phosphoproteomics and N-glycoproteomics**

Chin-Wen Chen[1], Pei-Yi Lin[1], Ying-Mi Lai[2], Miao-Hsia Lin[3], Shu-Yu Lin[4], Chuan-Chih Hsu[1,*]

[1]Institution of Plant and Microbial Biology, Academia Sinica, Taipei 115201, Taiwan
[2]Agricultural Biotechnology Research Center, Academia Sinica, Taipei 115201, Taiwan
[3]Department of Microbiology, National Taiwan University College of Medicine, Taipei 100233, Taiwan
[4]Academia Sinica Common Mass Spectrometry Facilities for Proteomics and Protein Modification Analysis, Academia Sinica, Taipei 115201, Taiwan

*corresponding author information:
Chuan-Chih Hsu


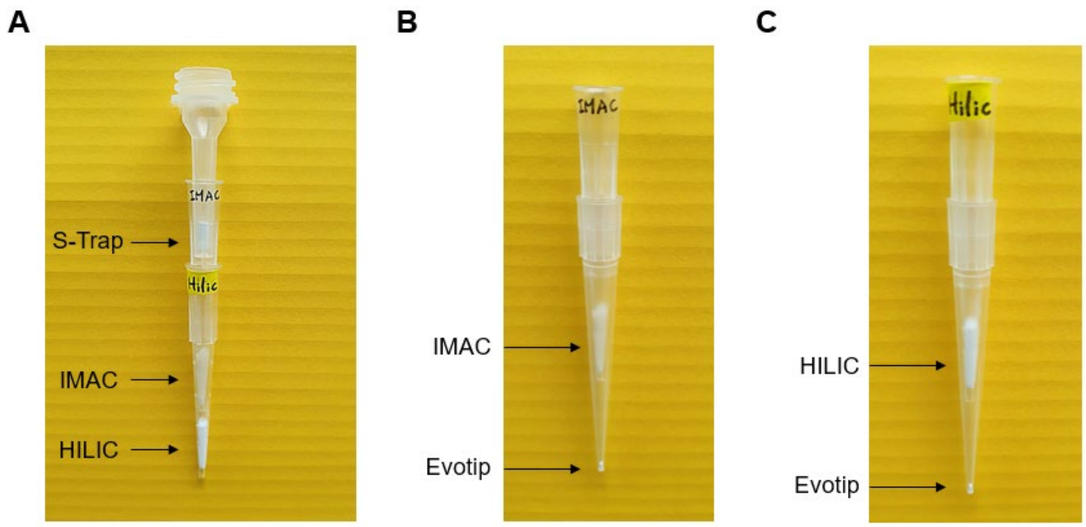


**Supplemental figure S1. Illustration of tandem tip strategies.** A, the tandem S-Trap-IMAC-HILIC strategy: The S-Trap micro column, IMAC tip, and HILIC tip are arranged in a tandem tip format and inserted into a tube for centrifugation. B, the tandem IMAC tip-Evotip arrangement. C, the tandem HILIC tip-Evotip arrangement.

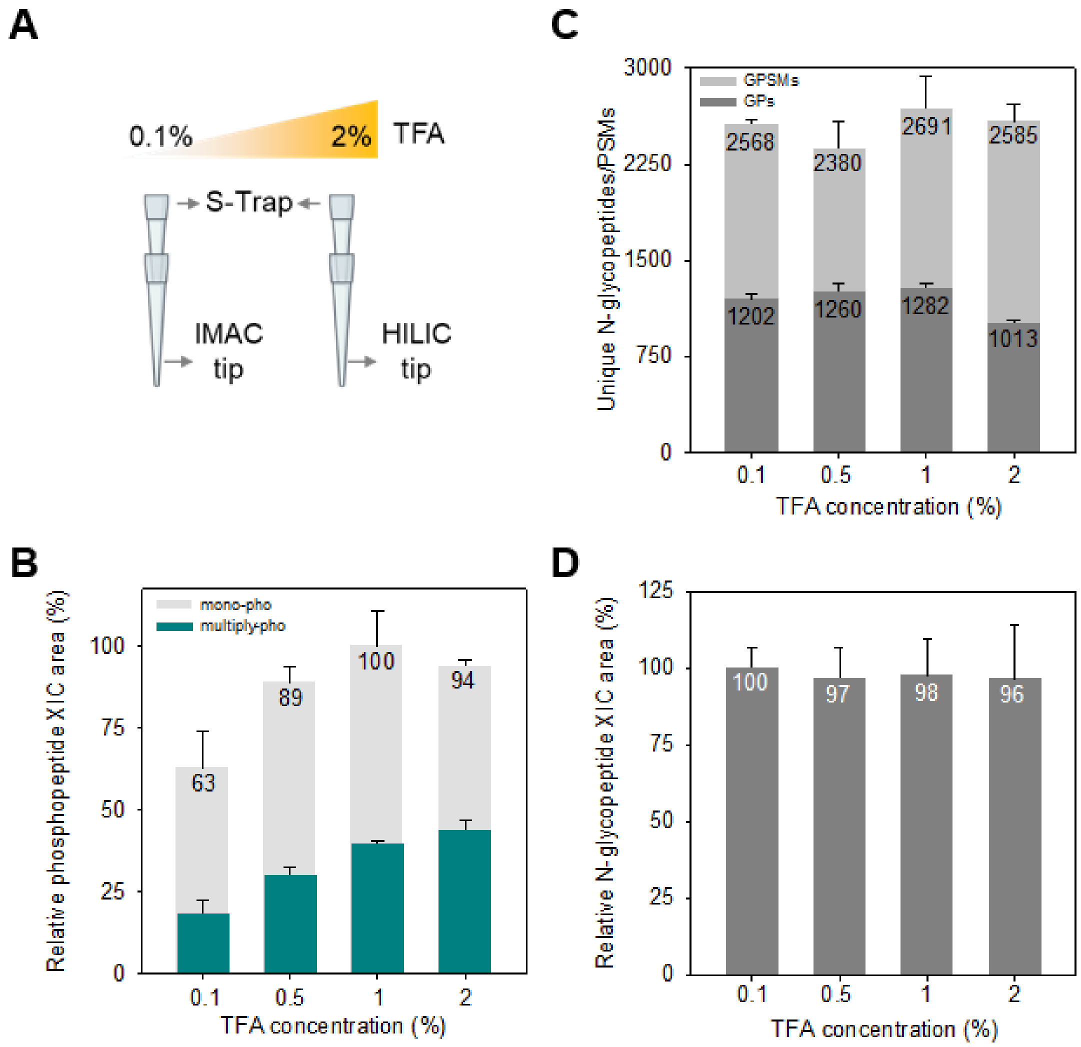


**Supplemental figure S2. Impact assessment of TFA concentration phosphopeptides and N-glycopeptides enrichment performance.** A, schematic of the experimental design for TFA concentration comparison. B, accumulated XIC area of identified phosphopeptides under four TFA concentration conditions (n = 3). Error bar, SD. C, number of identified GPSMs and GPs under four TFA concentration conditions (n = 3). Error bar, SD. Mono-pho, monophosphorylated peptide. Multiply-pho, multiply phosphorylated peptide. GPSMs, glycopeptide-spectrum matches. GPs, glycopeptides. D, accumulated XIC area of identified N-glycopeptides under four TFA concentration conditions (n = 3). Error bar, SD.

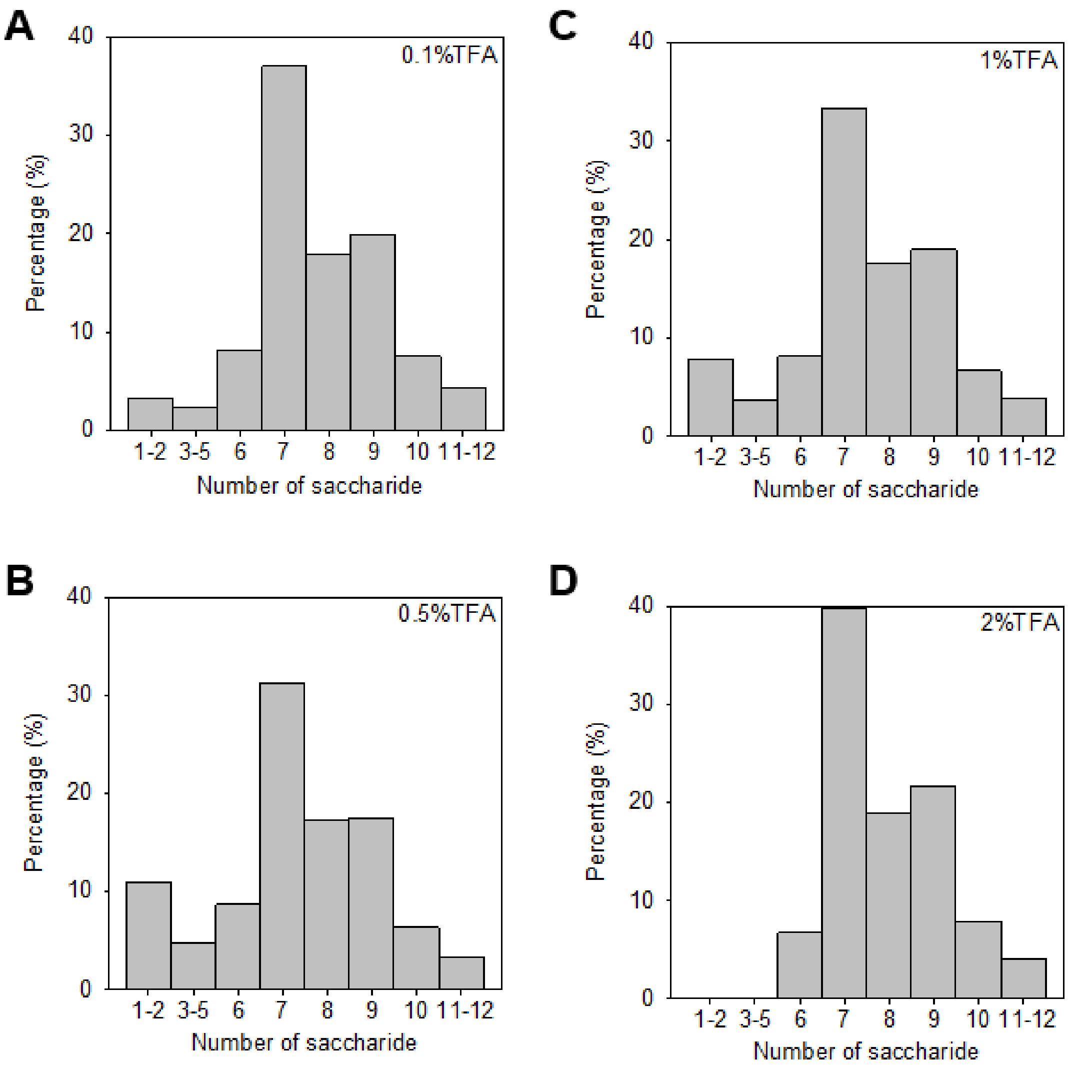


**Supplemental figure S3. Distribution of the number of saccharides within the glycan moieties of the identified N-glycopeptides at four TFA concentrations.** A, 0.1% TFA. B, 0.5% TFA. C, 1% TFA. D, 2% TFA.

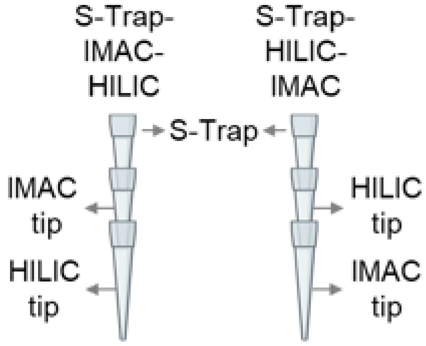


**Supplemental figure S4. Schematic representation of the S-Trap-IMAC-HILIC and S-Trap-HILIC-IMAC strategies designed for performance comparison.**

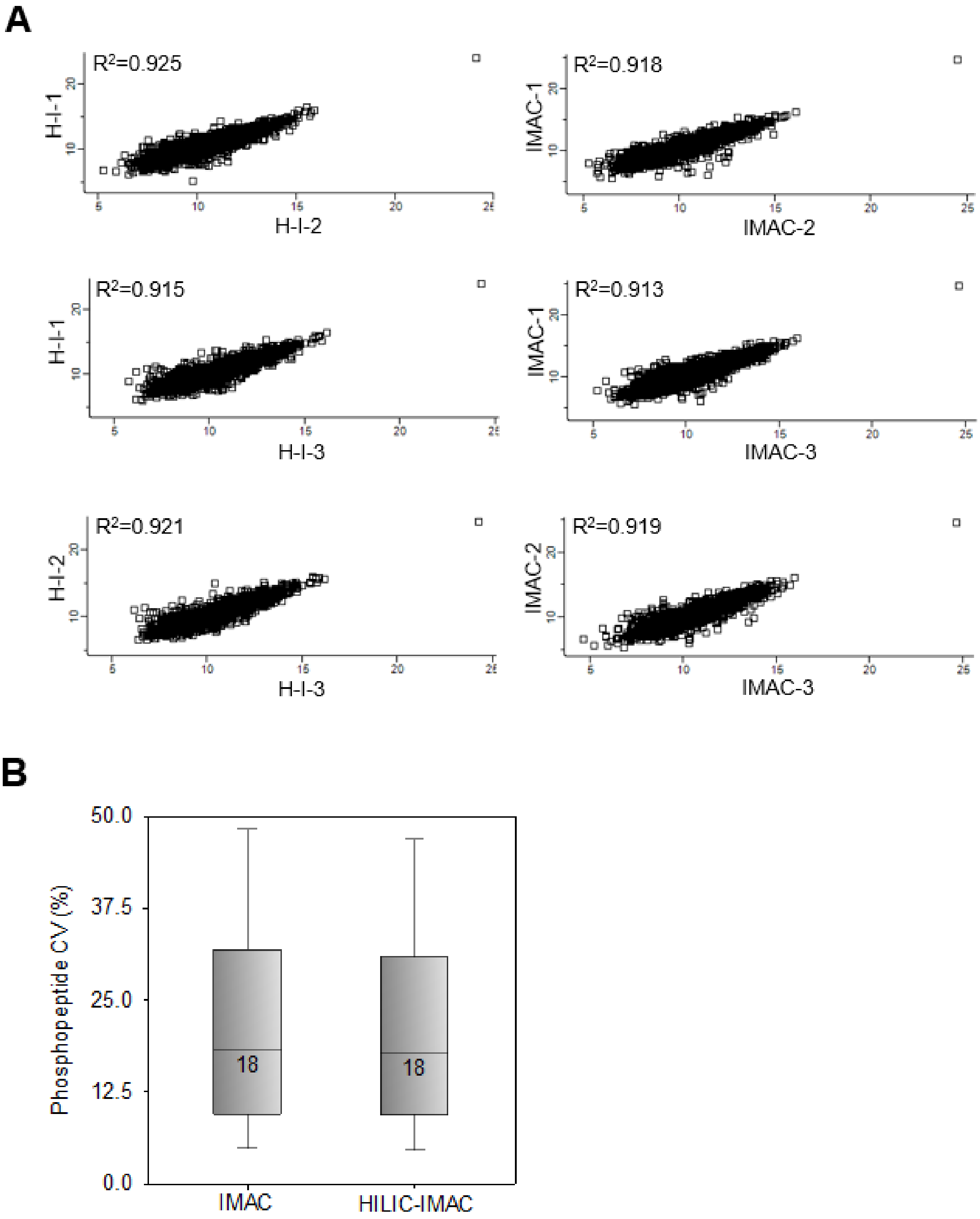


**Supplemental figure S5. Evaluation of reproducibility of the phosphopeptides identified in the IMAC and HILIC-IMAC protocols.** A, Pearson correlation analysis between technical replicates of the IMAC and HILIC-IMAC protocols. Log2-transformed phosphopeptide intensities were plotted on both the x- and y-axes. B, distribution of the log2-transformed phosphopeptide intensities identified from the IMAC and HILIC-IMAC protocols. Boxes mark the first, median, and third quantiles, and whiskers mark the minimum/maximum value within 1.5 interquartile range. "H-I" represents the HILIC-IMAC protocol.

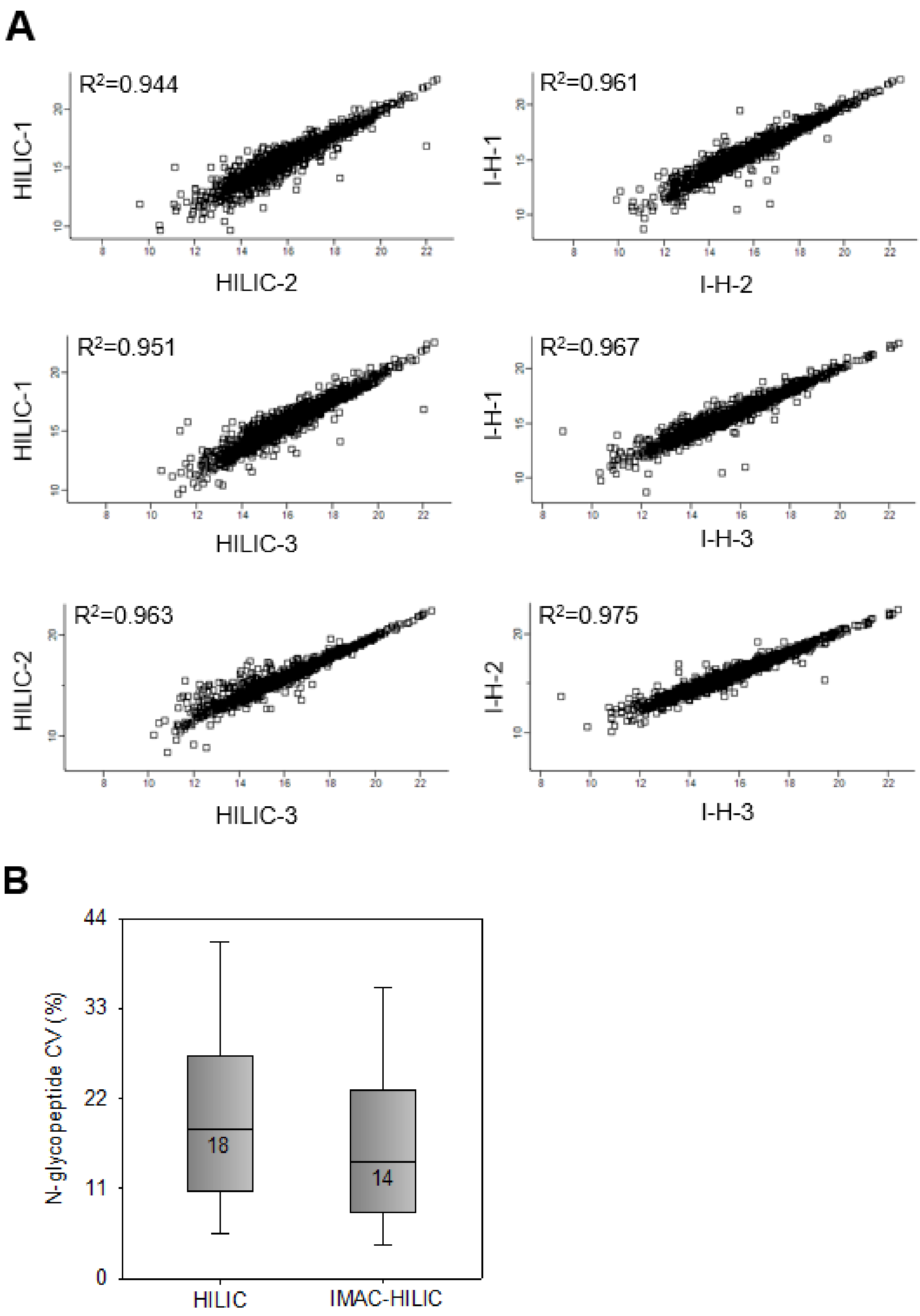


**Supplemental figure S6. Evaluation of robustness of the N-glycopeptides identified in the HILIC and IMAC-HILIC protocols.** A, Pearson correlation analysis between technical replicates of the HILIC and IMAC-HILIC protocols. Log2-transformed N-glycopeptides intensities were plotted on both the x- and y-axes. B, distribution of the log2-transformed N-glycopeptide intensities identified from the HILIC and IMAC-HILIC protocols. Boxes mark the first, median, and third quantiles, and whiskers mark the minimum/maximum value within 1.5 interquartile range. "I-H" represents the IMAC-HILIC protocol.

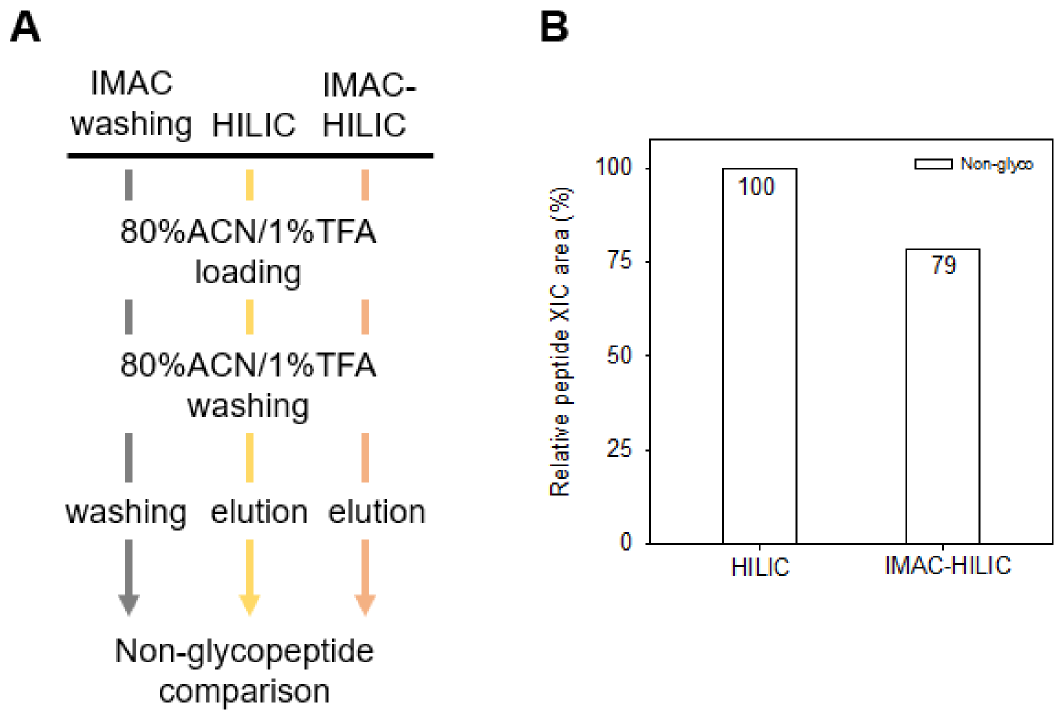


**Supplemental figure S7. Comparison of non-glycopeptides identified in IMAC washing, HILIC elution, and IMAC-HILIC elution.** A, schematic representation of the IMAC washing, HILIC, and IMAC-HILIC strategies designed for non-glycopeptide comparison. Non-glyco, non-glycopeptides. B, summed XIC area of identified non-glycopeptides in both the HILIC and IMAC-HILIC conditions. IMAC washing, the second step of IMAC washing.

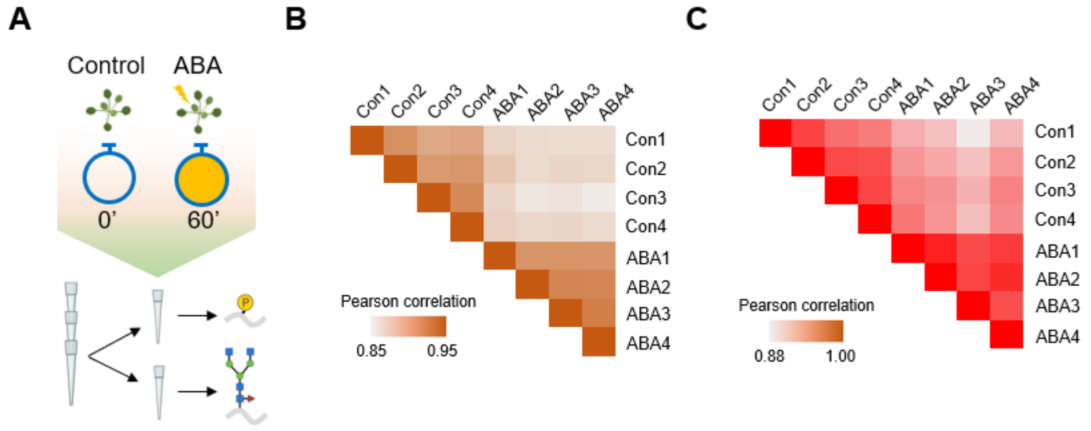


**Supplemental figure S8. Phosphoproteomics and N-glycoproteomics analysis of ABA-treated Arabidopsis seedlings.** A, experimental design for ABA-dependent phosphoproteomics and N-glycoproteomics analysis using the TIMAHAC approach. B, scatter plot shows the phosphoproteomics Pearson correlation of replicates in ABA-treated and untreated samples. C, scatter plot shows the N-glycoproteomics Pearson correlation of replicates in ABA-treated and untreated samples.

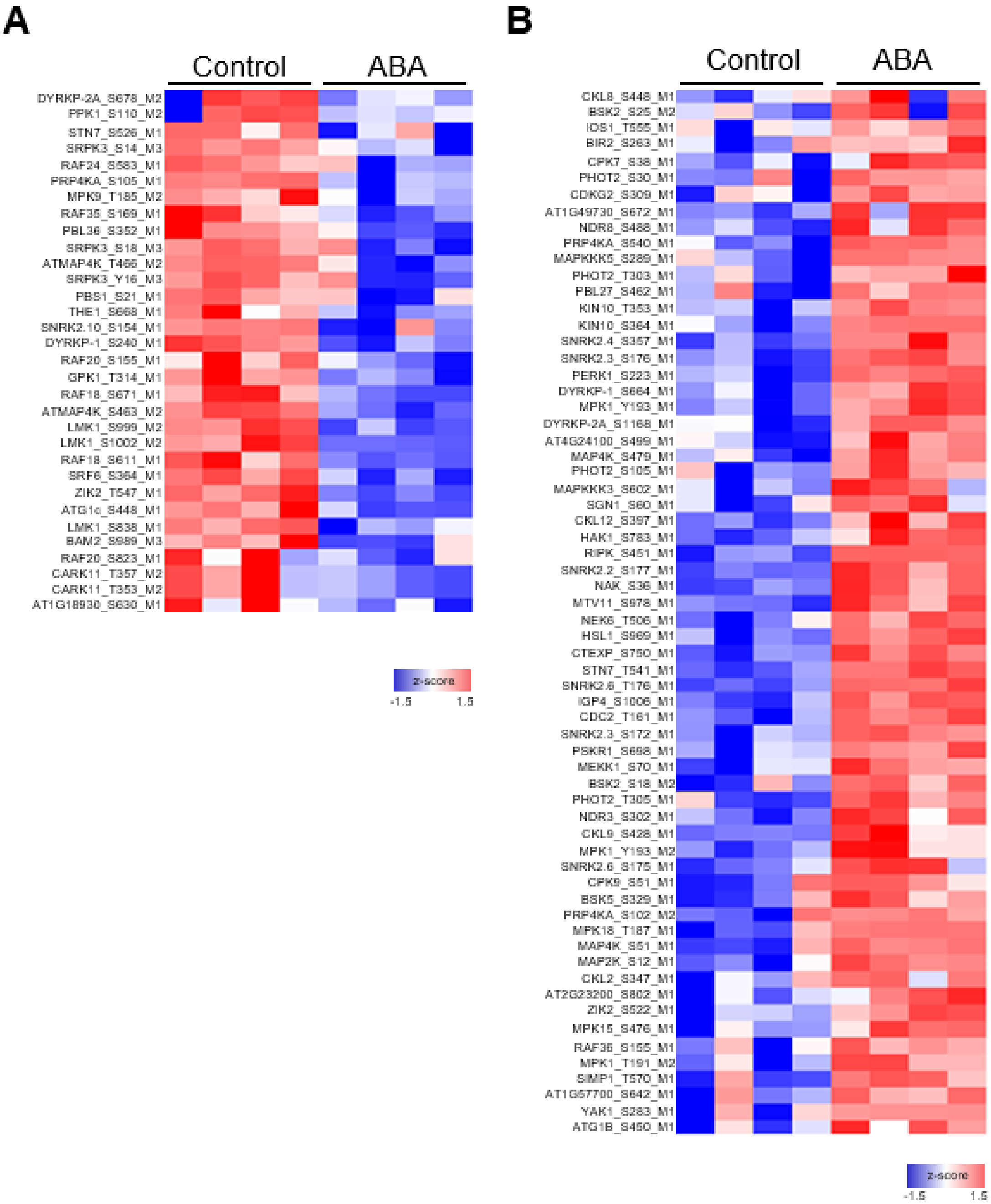


**Supplemental figure S9. Heatmaps of z-scored intensities of phosphorylation events on kinases significantly perturbated upon ABA treatment.** A, the ABA-repressed phosphorylation sites. B, the ABA-induced phosphorylation sites.

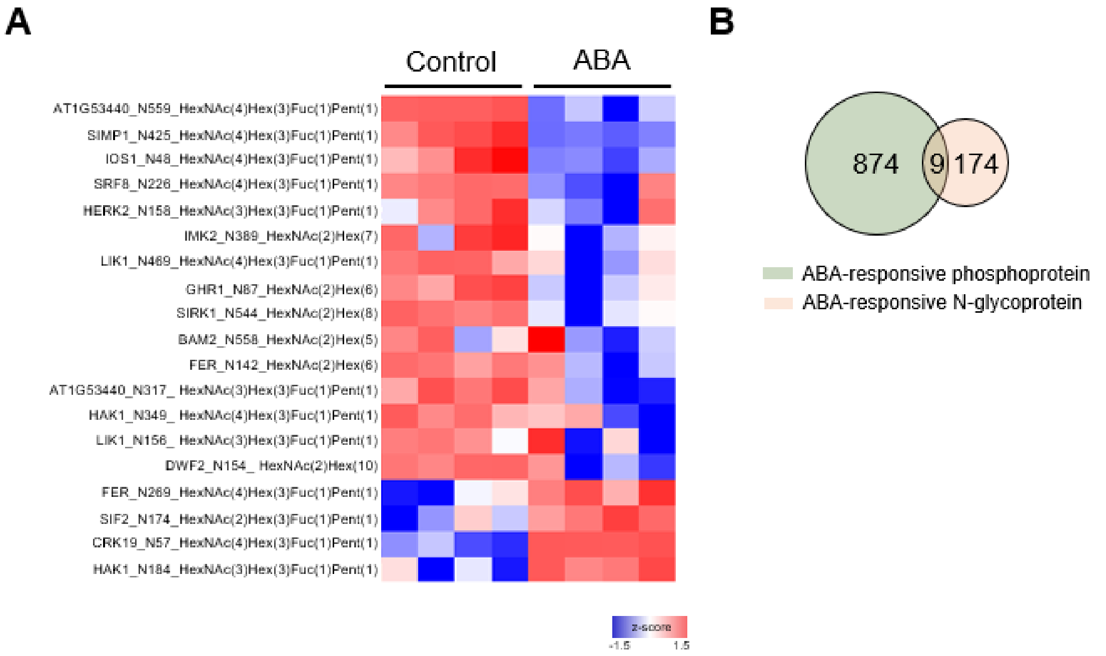


**Supplemental figure S10. Comparison of ABA-responsive phosphoproteins and N-glycoproteins.** A, heatmap of z-scored intensities of N-glycoforms on kinases significantly perturbated upon ABA treatment. B, Venn diagram representing the overlap of ABA-responsive phosphoproteins and N-glycoproteins.
